# *Drosophila* ovarian germline stem cell cytocensor projections dynamically receive and attenuate BMP signaling

**DOI:** 10.1101/370486

**Authors:** Scott G. Wilcockson, Hilary L. Ashe

## Abstract

In the *Drosophila* ovarian germline, BMP signals released by niche cells promote germline stem cell (GSC) maintenance. Although BMP signaling is known to repress expression of a key differentiation factor, it remains unclear whether BMP-responsive transcription also contributes positively to GSC identity. Here, we identify the GSC transcriptome using RNA-seq, including the BMP-induced transcriptional network. Based on these data, we provide evidence that GSCs form two types of cellular projections. Genetic manipulation and live *ex vivo* imaging reveal that both classes of projection allow GSCs to access a reservoir of Dpp held away from the GSC-niche interface. Moreover, microtubule-rich projections, termed ‘cytocensors’, form downstream of BMP and have additional functionality, which is to attenuate BMP signaling. In this way cytocensors allow dynamic modulation of signal transduction to facilitate differentiation following GSC division. This ability of cytocensors to attenuate the signaling response expands the repertoire of functions associated with signaling projections.

## INTRODUCTION

The stem cell niche is a tissue microenvironment, specialised in structure and function, that ensures the self-renewal and survival of cells needed to maintain tissue homeostasis throughout an organism’s life. The first niche was characterised in the *Drosophila* ovarian germline (Cox et al., 1998; King and Lin, 1999) where the Bone Morphogenetic Protein (BMP) family member, Decapentaplegic (Dpp), was found to be necessary for maintenance of germline stem cells (GSCs) (Xie and Spradling, 1998, 2000). Since this discovery there has been an explosion in the identification and characterisation of stem cell niches in most tissues and model organisms (Scadden, 2014).

Within the *Drosophila* ovary, GSCs are maintained at the anterior tip in discrete structures called germaria (Lin and Spradling, 1993). A small population of somatic cells, the Cap cells (CpCs), contact the GSCs through E-cadherin (Ecad)-based adherens junctions (AJs) (Song et al., 2002) and promote stem cell identity through the secretion of Dpp homodimers or Dpp-Glassbottom boat (Gbb) heterodimers. Dpp signals at exquisitely short-range to maintain 2-3 GSCs per niche. Upon cell division, one daughter cell exits the niche, allowing it to move out of the range of the Dpp signal and differentiate into a cystoblast (CB). Multiple mechanisms have been described for restricting Dpp range, including stabilisation or concentration of Dpp within the niche by the heparan sulphate proteoglycan (HSPG) Divisions abnormally delayed (Dally), sequestration by a collagen IV matrix in between the GSCs and CpCs, and escort cell (EC) expression of the Dpp receptor, Thickveins (Tkv), which acts as a ‘decoy’ to soak up any free BMP ligand (Wilcockson et al., 2017). The most anterior ECs thus define the posterior limit of the GSC niche and contact the differentiating CBs to create a differentiation niche.

Within GSCs, the BMP signal is transduced by phosphorylation and activation of the Smad1/5 ortholog, Mothers against Dpp (Mad). Mad oligomerises with the Smad4 ortholog Medea leading to their nuclear accumulation (Hamaratoglu et al., 2014). A key Dpp target gene in GSCs is *bag of marbles* (*bam*), encoding an essential differentiation factor, which is repressed by Dpp signaling (Chen and McKearin, 2003). Upon cell division, the daughter cell that exits the niche derepresses *bam*, which initiates the differentiation programme. However, few Dpp target genes have been identified in GSCs and there is little understanding of how the BMP self-renewal signal may positively act on GSC identity. Therefore, we used RNA-seq to identify the GSC transcriptional network, including genes that are regulated by the Dpp signal. These data reveal that the GSC synthesises different types of cellular projections that function to receive the niche BMP signal, including one class that also plays an active role in BMP signal attenuation, which we thus refer to as ‘cytocensors’.

## RESULTS

### RNA-seq of GSC-like cells and CBs reveal putative GSC self-renewal and maintenance factors

In order to identify regulators of GSC self-renewal and differentiation, we compared the GSC and CB transcriptomes, by purifying these cells based on the expression of known cellular markers. Expression patterns and further information on the cell types and signaling circuitry in the germarium are shown in Fig. S1A-F. In the absence of a GSC-specific marker, we genetically expanded the GSC population by expressing constitutively active Tkv (*UASp-tkv^QD^*) using a maternal germline *Gal4* driver (*nos-Gal4::VP16*) in a *vasa^GFP^* background. Vasa is a germ cell marker that we used to isolate GSCs by FACS (Fig. 1A, Sano, et al., 2002). Flies of this genotype form tumours of pMad^+^ GSC-like cells identifiable by a single, round spectrosome, a germline-specific spectrin-rich endomembrane organelle that becomes branched in more developed cysts (Fig. S1G). CBs were isolated by FACS based on their expression of a *bam*-*GFP* reporter and as single cells to exclude more developed GFP^+^ cysts (Fig. S1E, Chen & Mckearin, 2003). Differential expression analysis revealed 2,249 differentially expressed genes with around one third up-regulated in *tkv^QD^* (GSCs) and two thirds up-regulated in *bam.GFP* expressing cells (CBs) (Fig. 1A, Table S1), including *bam*. GO term analysis of GSC- and CB-enriched transcripts reveals distinct biological processes (Fig. 1B), including nervous system development and cell migration for GSCs. Enriched transcripts within these categories encode adhesion proteins, axon guidance molecules, ciliogenesis factors and structural/cytoskeletal proteins.

**Figure 1.**
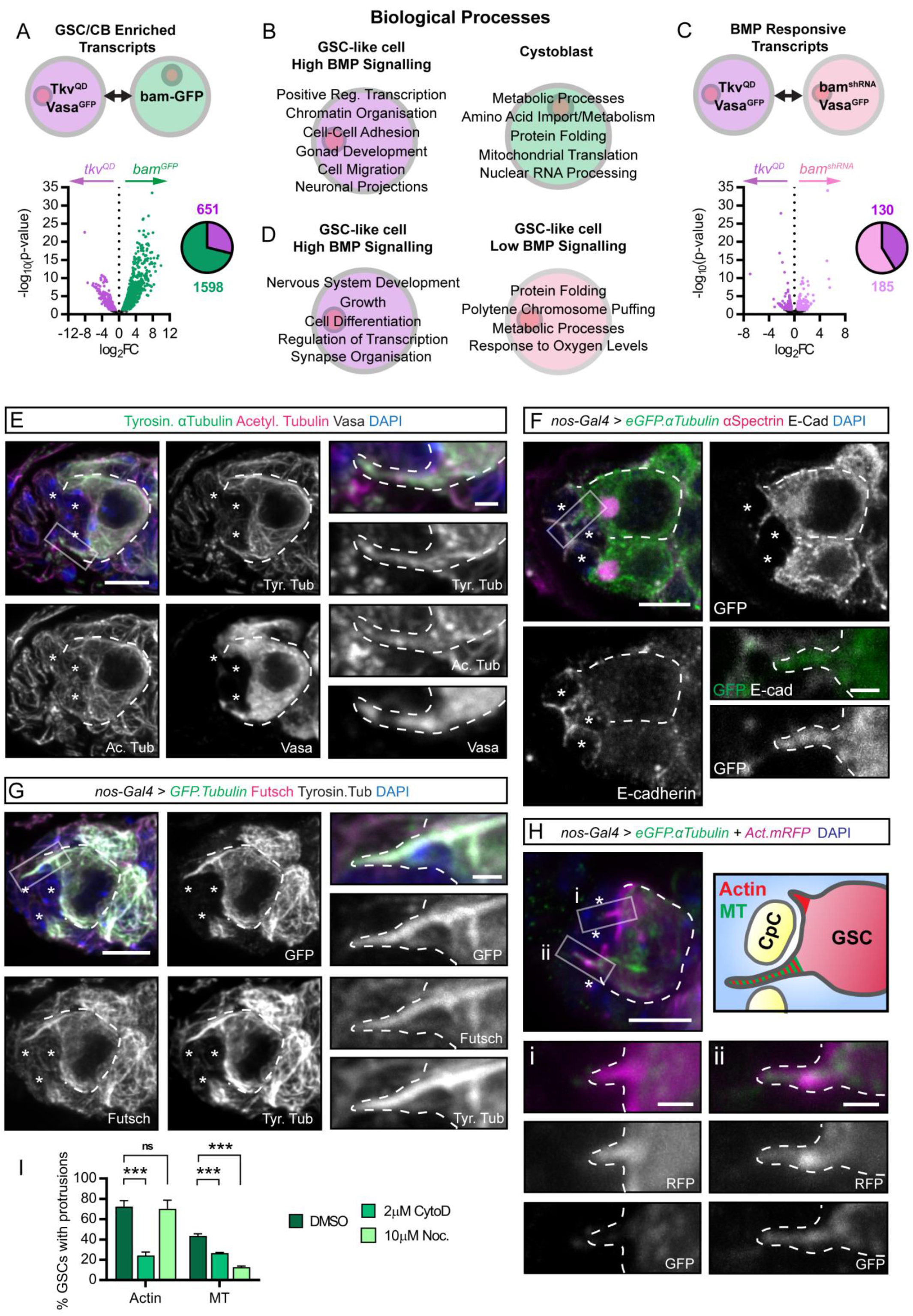
GSCs up-regulate MT-associated factors and extend cytoskeletal projections into the niche. (A) Differential expression analysis of RNA enriched in *tkv^QD^* (magenta) and *bam.GFP* (green) expressing GSC-like cells and CBs, respectively. Pie chart shows number of significantly enriched genes for each cell type (log_2_Fold change>0.5, p<0.05). (B) GO term analysis results showing biological processes enriched in in *tkv^QD^* (magenta) and *bam.GFP* (green) expressing cells. (C-D), as in (A-B) comparing *tkv^QD^* (magenta) and *bam^KD^* (light pink) expressing GSC-like cells. (E) The MT network of the germarium. GSCs are marked by Vasa expression (white). MTs are labelled by the MT markers; acetylated αTubulin and tyrosinated αTubulin. (Inset) closeup views of the indicated boxed region showing a GSC MT-rich cytoplasmic projection. (F-G) Immunofluorescence staining of germaria with germline *eGFP.αTub* expression. GSCs marked by the spectrosome visualised with anti-αSpectrin, Ecad outlines the CpCs (*) in F. (G) The MT-associated factor Futsch also localises to GSC MT-rich projections. (Inset) closeup views of the indicated boxed regions. (H) GSCs form different actin-rich projections. Immunofluorescence staining of germaria with germline expression of *eGFP.αTub* and *Act.mRFP*. A GSC extends one MT- and actin-rich projection (i) and one actin-based filopodium (ii). (I) Percentage of GSCs forming projections after 30min *ex vivo* treatment of *nos-Gal4>Act42A.GFP or eGFP.αTub* ovaries with DMSO (control), 2µM CytoD or 10µM nocodazole. Mean and SD from n>100 GSCs; n=3 biological replicates. ns, not significant; Dashed lines outline individual GSCs and (insets) projections. (*) CpCs. Scale bar = 5µm or 1µm (insets). ***, p<0.0001. See also Figure S1 and S2.

Having identified the GSC transcriptome, we next identified the subset of genes specifically regulated by Dpp signaling, by using germline-specific RNAi knockdown of *bam* expression (*UASp-bam^KD^*) which blocks GSC differentiation. Germ cells that exit the niche continue to divide away from the short-range Dpp signal and therefore form tumours of pMad^-^ GSC-like cells with a single, round spectrosome, that can again be isolated through *vasa^GFP^* expression (Fig. S1H). Differential expression analysis of *tkv^QD^-* and *bam^KD^*-expressing GSC-like cells allows the comparison of ‘high Dpp’ and ‘low Dpp’ GSCs, respectively. This reveals around 300 genes differentially regulated by Dpp signaling, with just under half the genes up-regulated in response to Dpp (Fig. 1C, Table S2), including *dad*. GO term analysis identifies processes activated and repressed by Dpp signaling (Fig. 1D). Again, genes upregulated in response to Dpp signaling encode proteins involved in nervous system development and synapse organisation, including the master ciliogenesis transcription factor *Rfx* and the microtubule (MT)-associated protein 1B (MAP1B) homolog *Futsch*. Together these data define the early germline transcriptome, from self-renewing GSCs to differentiating daughter CBs, and the subset of this network functioning downstream of Dpp signaling.

### Germline stem cells extend MT- and actin-rich projections into the niche

Putative Dpp target genes in the GSC transcriptome include *Rfx* and *futsch*, which both regulate the formation of MT-based structures. To investigate a potential cytoskeletal response to Dpp signaling, we first defined the stem cell MT network using immunofluorescence staining of tyrosinated α-tubulin (a marker of new, dynamic MTs), acetylated α-tubulin (a marker of stable MTs), and the germ cell marker, Vasa. GSCs are enriched for tyrosinated α-tubulin compared to the post-mitotic CpCs (Fig. 1E, CpCs marked by asterisks and the GSC is outlined by a dashed line). This difference enables the visualisation of stem-derived MT-rich projections that a subset of GSCs extend into the niche (Fig. 1E, see box). The cytoplasmic protein, Vasa, also localises within these projections, confirming these structures are GSC-derived. To further characterise these MT-rich projections, we specifically visualised the stem cell MT network through the germline expression of *UASp*-*eGFP.αTubulin84B* (*GFP.αTub*). Ecad staining delineates the contact points between individual CpCs and the GSC-niche interface. A subset of GSCs is found to generate a single, short MT-based projection that extends around or between the CpCs (Fig. 1F, S2A). These data reveal that ovarian GSCs generate MT-based structures that extend toward and between the niche CpCs.

As our RNA-seq data identified genes associated with ciliogenesis we addressed whether these MT-rich projections were ciliary in nature. GFP.αTub^+^ projections are composed of acetylated tubulin, a classical ciliary marker (Fig. S2B, Bi). However, the MTs show a non-uniform pattern of acetylation unlike stable ciliary MTs. In addition, no association of these GSC projections with the centrosome, based on γ-tubulin staining, is observed (Fig. S1C). Therefore, we conclude that these MT-rich projections are not ciliary in nature and hereon refer to them as ‘cytocensors’ based on their signal suppression property, ie acting as a censor (see later). GSCs of the *Drosophila* testis generate MT-nanotubes whose formation is regulated by ciliary proteins (Inaba et al., 2015). However, our data highlight numerous differences between cytocensors and MT-nanotubes, consistent with them being distinct structures (see Discussion).

One of the positive Dpp target genes identified in the RNA-seq is *futsch*. Futsch function is best characterised in the nervous system where it promotes MT stability (Halpain and Dehmelt, 2006). Futsch staining reveals strong GSC enrichment compared with CpCs (Fig. 1G). Futsch co-localises extensively with the stem cell MT network, including the cytocensor MTs. These data are consistent with Dpp signaling up-regulating *futsch* expression and Futsch subsequently localising to GSC cytocensors where it may play a role in projection stability (see later).

We next addressed whether these projections contain or require the actin cytoskeleton for their formation by specifically expressing both *GFP*.*αTub* and *UASp-Actin5C.mRFP* in the germline. Immunofluorescence staining reveals that GSCs also generate multiple actin-based projections (APs). Figure 1H shows a single stem cell with two distinct projections; one short actin-based filopodium (Fig. 1H box i) and a longer projection containing both MTs and actin (Fig. 1H box ii). This shows that GSC cytocensors are both MT- and actin-rich projections, while GSCs also generate additional APs.

Having identified that ovarian GSCs extend multiple distinct cellular projections, we investigated the regulatory relationship between actin and MTs in the formation of these projections. We treated *Drosophila* ovaries *ex vivo* with cytochalasin D (CytoD) and nocodazole, inhibitors of actin and MT polymerisation, respectively. A short 30 min incubation with CytoD or nocodazole was sufficient to disrupt F-actin (Figure S2D) or reduce tubulin levels (Fig. S2E), respectively. Treatment of ovaries expressing germline *UASp-Act42D.GFP* with CytoD significantly reduces the number of niche-directed APs while nocodazole had no effect (Fig. 1I). Conversely, treatment of *GFP.αTub* expressing ovaries with both drugs resulted in a significant reduction in the ability of GSCs to generate cytocensors, in comparison with DMSO-treated ovaries. This suggests that APs form independently of the MT network, however, the formation of cytocensors is dependent on actin. In addition, APs are more abundant than cytocensors (Fig. 1I). The relative abundance of APs and the requirement for actin polymerisation for cytocensor formation suggests that APs may represent the primary structure from which cytocensors emanate.

### Projections dynamically probe the niche microenvironment

The above data showing enrichment of tyrosinated MTs and the non-uniformity of α-tubulin acetylation suggests that the cytocensors may be dynamic in nature. To address this, we used live imaging to monitor MT dynamics in *GFP.αTub* expressing GSCs *ex vivo*. GSCs extend cytocensors that dynamically probe the niche microenvironment over the course of 1-2 hours (Video S1). Figure 2A shows stills from Video S1 that focuses on a single motile cytocensor extended into the niche that collapses and reforms multiple times over the course of imaging. This shows that cytocensors are relatively dynamic in nature. We similarly visualised F-actin by driving germline expression of *UASp-LifeAct.eGFP* (Video S2). Strong labelling of cortical F-actin is seen at the GSC-niche interface from which the APs emerge (Fig. 2B). The GSCs generate short filopodia-like projections that extend into the niche and although they appear dynamic, a subset exhibit much longer lifetimes (compare the relatively short-lived AP in Fig. 2Bi with a longer-lived one in Bii). We also found additional transient lateral GSC projections (Fig. 2C, magenta arrowhead, Video S3) while all early differentiating germ cells (CBs (green), 2- to 4-cell cysts (blue)) generate long, transient APs (Fig. 2Ci, ii). In addition, some GSCs extend broad lamellipodia-like projections that extend over or in between multiple niche cells (Fig. 2Ci-iii, Video S3), and individual finger-like filopodia extend from the ends of these structures to envelope niche CpCs (Fig. S3A, Video S4). During mitosis GSCs undergo typical cell rounding, accompanied by cortical F-actin accumulation and the collapse of APs, which rapidly reform as the GSCs re-establish contact with niche CpCs (Fig. S3B, Video 5).

**Figure 2.**
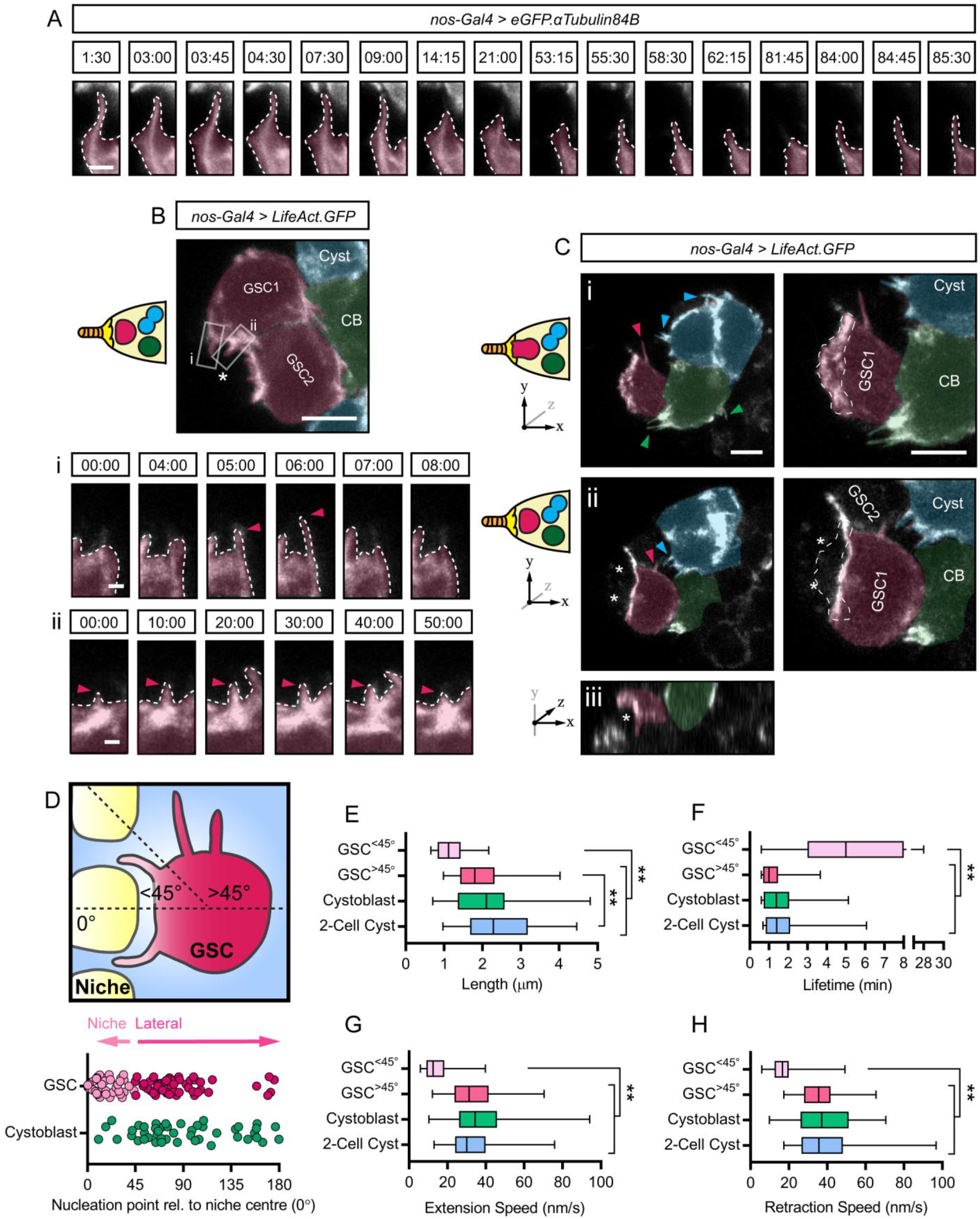
GSC projections are dynamic and all early germ cells form actin-rich projections. (A) Stills from Video 1 showing a cytocensor labelled with *eGFP.αTub* (false coloured magenta) (Time in mins). Image only shows part of the GSC to enable visualisation of the projection. (B) Stills from Video 2 showing F-actin with *LifeAct.GFP* in GSCs (false coloured magenta), CBs (green) and 2- or 4-cell cysts (blue). (Bi and ii) show stills of the regions in indicated boxes (Time in mins). Arrowheads indicate tip of the filopodium. (C) same as (B) showing stills from Video 3. (i, left) The first two *z-*slices of a maximum projection showing the formation of F-actin-rich projections by GSCs, CBs, and cysts, indicated by boxes and colour-coded arrowheads. (i, right) A closer view of the GSC in (i, left) showing a broad lamellipodia-like projection (outlined by dashed line) depicted in cartoon form on the left and axes denote position within the maximum projection. (ii, left) Two *z-*slices in the middle of the maximum projection reveal two CpCs (*) positioned below the lamellipodium. (ii, right) A closer view of the GSC in (ii, left) with the position of the overlying lamellipodium indicated by the dashed line. (iii) *xz*-plane view. (D) Cartoon and scatter plot showing the angle of actin filopodia nucleation point relative to the centre of the niche (0°). ‘Niche-directed’ filopodia are defined as those forming at an angle <45° to the centre of the niche. The rest are ‘lateral projections’. (n=100 GSC projections and n=50 CB projections) (E-I) Box and whisker plots comparing the length (E), lifetime (G), extension (H) and retraction speed (I) of actin projections from the following classes; GSC^<45°^ (light pink), GSC^>45^ (magenta), CBs (green) and 2-cell cysts (blue). Median, 25^th^ and 75th percentile and whiskers show minima and maxima. n=46- 50 projections from n≥8 cells each. Dashed lines outline GSC projections. Scale bar = 2µm (A) or 5µm (B-C). ***, p<0.0001. See also Figure S3.

GSCs appear to extend shorter projections toward the niche with lateral projections tending to be longer and more transient (Fig. 2C, Video S3). To compare the nature of these projections, we grouped together those that extend into the niche (GSC^<45°^; point of nucleation occurs at <45° relative to the centre of the niche (0°)) and those that extend laterally (GSC^>45°^; point of nucleation occurs at >45° is defined as a lateral projection) and then compared these to the projections generated by the differentiating cells (Fig. 2D). GSC projection formation appears polarised as most APs are nucleated in the direction of the niche, or below 90°, while CBs tend to generate projections at any angle >45° (Fig. 2D). This may simply be due to structural hinderance, with the presence of neighbouring GSCs precluding the formation of niche-directed CB projections. Comparing the length, lifetime, and speed of filopodia extension and retraction reveals that GSC^<45°^ projections are significantly shorter, slower and more stable than GSC^>45°^ projections (Fig. 2E-H). These lateral projections appear similar to those formed by CBs and 2-cell cysts; they are generally longer, the extent of which increases as differentiation progresses (Fig. 2E), and significantly more transient/unstable, indicated by their short lifetime (Fig. 2F) and their speed of growth and collapse (Fig. 2G-H). Together these data show that GSC projections are dynamic and probe the niche. In addition, the formation of unstable APs is a common trait of early germ cells, while GSCs also extend more stable, short APs into the niche.

### Cytocensors form in response to Dpp signaling

Our RNA-seq data suggest that Dpp signaling activates *Futsch* and *Rfx* expression, while GSCs also showed general enrichment of other cytoskeletal- and ciliogenesis-associated factors. We therefore determined whether a subset of these factors plays a role in the formation of GSC cytocensors. To achieve this, germline-specific RNAi was used to firstly address the roles of the MT-stabiliser Futsch and the tubulin-binding protein Stathmin (Stai). *futsch^KD^* expression significantly reduces the frequency of cytocensors formed, while those that are formed are shorter than wildtype projections (Fig. 3A-D). Conversely, *stai^KD^* expression leads to significantly longer and thicker projections. We also knocked down expression of Klp10A, a MT-depolymerising kinesin shown to regulate MT-nanotube formation and centrosome size in male GSCs (Chen et al., 2016; Inaba et al., 2015). This also leads to significantly longer and slightly thicker projections than wildtype (Fig. 3A-D). Klp10A does not however appear to regulate female GSC centrosome size (Figure S4A-Ai).

**Figure 3.**
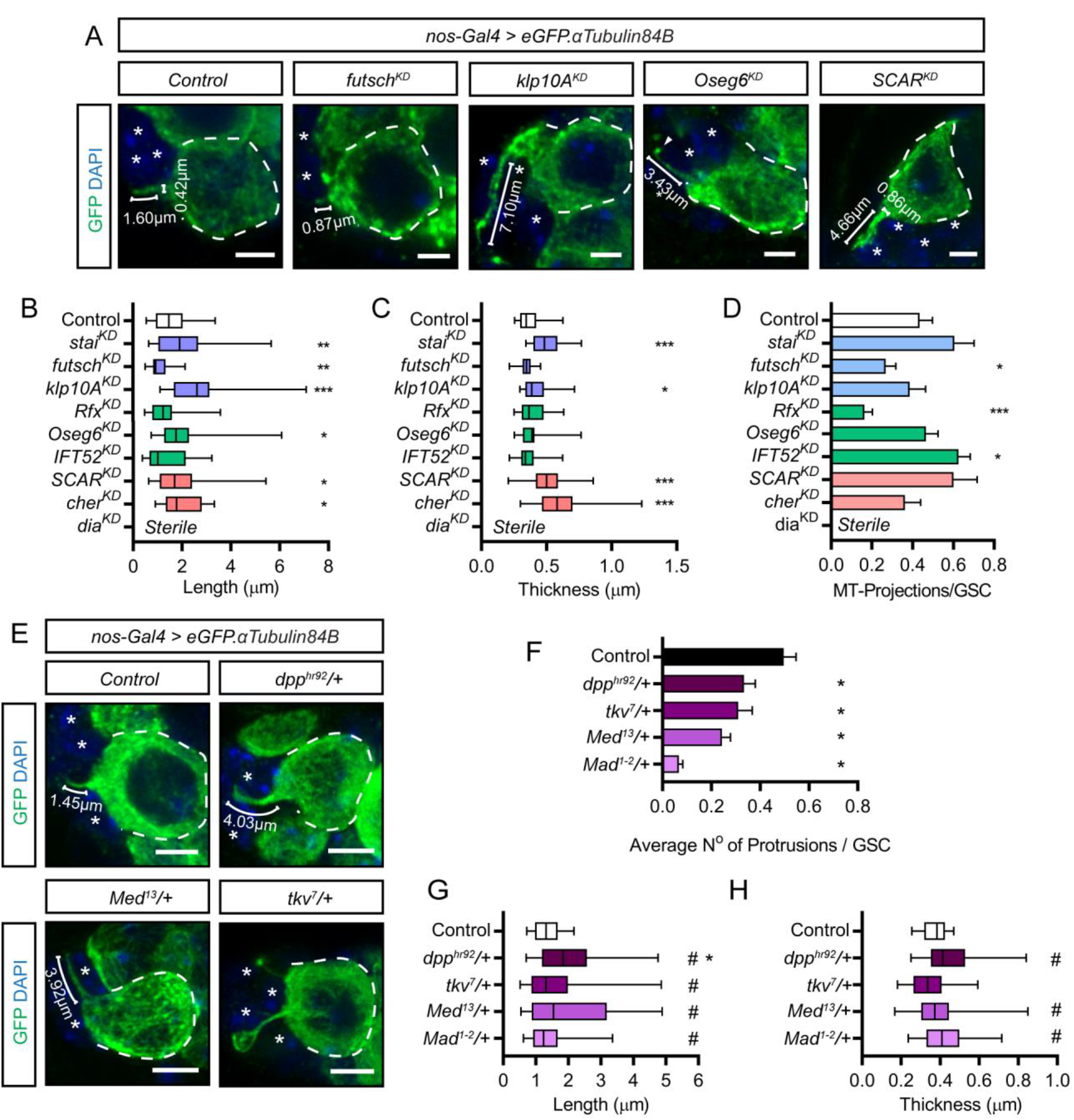
Cytocensors are regulated by stem cell enriched MT-associated factors and form in response to Dpp signaling. (A) Germline-specific *shRNA e*xpression for the indicated MT-associated factors and actin regulators disrupts projection formation. (B-C) Box and whisker plots of cytocensor length (B) and thickness (C) following knock down of the expression of MT-associated factors (blue), ciliogenesis factors (green) and actin regulators (red). Median, 25^th^ and 75th percentile and whiskers show minima and maxima. n=46-50 projections from n>8 cells. (D) Bar chart showing number of cytocensors formed per GSC for knockdown of factors as in (B-C). Mean and SEM. (E-H) Heterozygous mutants for Dpp signaling pathway components, *dpp^hr92^*, *tkv^7^*, *med^13^* and *mad^1-2^*, typically form abnormal cytocensors compared to controls. Reduced Dpp signaling disrupts projection formation (F), length (G) and/or thickness (H). Statistics as in (B-D). #, p<0.01 F-test. Dashed lines outline individual GSCs. (*) CpCs; brackets, length or thickness. Scale bars = 2µm. *, p<0.05; **, p<0.001, ***, p<0.0001. See also Figure S4.

We next addressed the roles of three genes associated with ciliogenesis. *Rfx^KD^* expression leads to a significant decrease in the frequency of cytocensor formation (Fig. 3D), while those that are formed appear normal in length and thickness (Fig. 3B-C). Two intraflagellar transport proteins were also included in our analysis, Oseg6 and IFT52, which are expressed in GSCs and CBs (Table S1) and function downstream of Rfx (Laurençon et al., 2007). *Oseg6^KD^* expression results in longer cytocensors that can often be found to contain a globular accumulation of tubulin at the tip (Fig. 3A), while *IFT52^KD^* expression resulted in a small increase in the frequency of cytocensor formation. Together, these data identify a set of factors, two of which are up-regulated by Dpp signaling (Futsch and Rfx), that regulate cytocensor formation.

As actin is necessary for the formation of MT-rich cytocensors (Fig. 1I), we examined three actin cytoskeletal components and regulators of filopodia and cytoneme formation; the formin diaphanous (dia), SCAR, and filamin (cheerio, cher). *dia^KD^* expression results in complete loss of the germline, therefore the effect on cytocensor formation could not be determined. However, knockdown of either *SCAR* or *cher* expression resulted in the formation of abnormally long and thick projections (Fig. 3A-C). These results further highlight the role of actin in the formation of cytocensors.

Two of the genes validated as regulators of cytocensor formation are *Rfx* and *futsch*, both of which were identified as positive Dpp targets in our RNA-seq data. This suggests that cytocensor formation is downstream of Dpp signaling in GSCs. To test the requirement for Dpp signaling, we visualised cytocensor formation in germaria from female flies heterozygous for mutations of the ligand, receptor (*tkv*), or Smads (*mad* and *med*). Heterozygotes were used as a sensitised background because analysis of homozygous mutants is not possible due to the rapid differentiation of GSCs in the absence of Dpp signaling. All heterozygotes show a significantly reduced ability to form projections, with *mad^1-2^*/*+* causing the greatest loss (Fig. 3E-F). In addition, cytocensors that are formed are frequently abnormal, particularly longer and thicker than controls (Fig. 3G-H). These results show that the frequency of cytocensor formation correlates with the ability of GSCs to receive and transduce Dpp signaling.

To further test the role of Dpp signaling in cytocensor formation, we exploited the previous observation that knockdown of ColIV expression in larval haemocytes, which deposit ColIV within the niche, results in increased Dpp signaling range and the accumulation of ectopic GSCs outside the niche (Van De Bor et al., 2015). Haemocytes deposit niche ColIV whilst contributing only little ColIV to the rest of the ovary. We therefore utilised this specificity to address whether ectopic Dpp could induce cytocensor formation. In a wildtype germarium cells that exit the niche, and therefore do not receive Dpp, do not typically generate cytosensors (Fig. S4B). Prevention of ColIV deposition in the niche by larval haemocytes (*HmlΔ-Gal4* > *ColIV^KD^*) extends Dpp signaling range, resulting in the accumulation of ectopic GSC-like cells (Van De Bor et al., 2015) that extend cytocensors (Fig. S4C). These data are consistent with Dpp signaling acting as a regulatory input for cytocensor formation.

### CpC-presentation of Dally generates a reservoir of Dpp

To investigate the function of GSC niche directed projections we examined the localisation of Dpp around the niche using endogenous Dpp tagged with mCherry (Dpp^mCh^; Fereres et al., 2018), in the same position as the previously described Dpp^GFP^ (Entchev et al., 2000). These Dpp^mCh^ flies are homozygous viable and show no overt germarium phenotype (Fig. S5A-B). Using extracellular staining of Dpp^mCh^ and Ecad, which outlines CpCs, we detect Dpp^mCh^ concentrated in puncta to the anterior of the niche creating an anterior to posterior high to low gradient (Fig. 4A-Ai). This same localisation pattern is also observed using two previously described Dpp transgenic lines tagged with either HA (Fig. S5C, Shimmi et al, 2005) or GFP (Fig. S5D, Teleman & Cohen, 2000), which tag both forms of Dpp generated by preprotein cleavage. This shows that secreted Dpp is concentrated away from GSCs, creating a short anteroposterior gradient across the niche.

**Figure 4.**
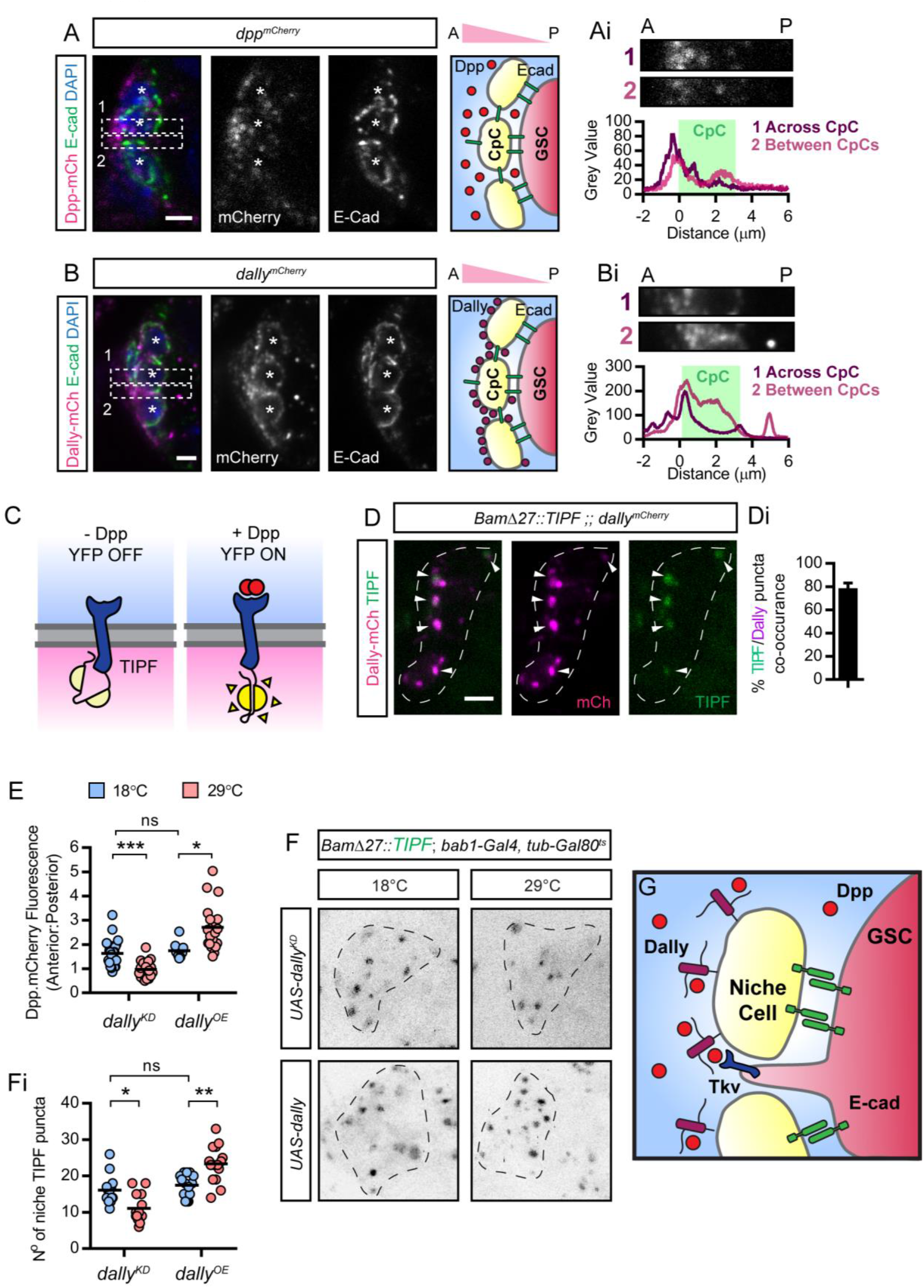
Niche cells create a Dally-Dpp reservoir where GSC Tkv activation occurs. (A-B) Extracellular staining of Ecad and endogenous mCherry-tagged Dpp (A) or Dally (B). Boxes show where the graphs of fluorescence intensity (Ai-Bi) were taken from anterior to posterior (A to P) through the centre of the niche or between two CpCs (*) as shown in the higher magnification views above. Ecad defines the niche cell boundaries (green). (C) Cartoon illustrating the TIPF reporter that fluoresces only upon ligand-receptor binding. (D) Endogenous fluorescence of Dally^mCh^ and germline expressed TIPF reporter (*BamΔ27::TIPF*) around the niche. (Di) Percentage co-occurrence of TIPF and Dally^mCh^ puncta. Mean and SD. n=20 germaria. (E) Graph shows anterior to posterior ratio of fluorescence intensity of Dpp^mCh^. *dally^KD^* or over-expression is induced at 29°C and compared to non-induced controls raised at 18°C. Line shows mean. n=20 CpCs. (F) Endogenous fluorescence of TIPF reporter with *dally^KD^* or over-expression in inverted black and white for clarity. (Fi) Graph showing the number of TIPF puncta per germaria in (F). Line shows mean, n=20 germaria. (G) Cartoon model showing that GSCs present Tkv on projections to access a Dally-Dpp reservoir. Dashed lines (D and G) outline the niche. Scale bar = 2µm. *, p<0.01; **, p<0.001; ***, p<0.0001. See also Figure S5.

The HSPG Dally is expressed by CpCs and promotes Dpp signaling in GSCs (Guo and Wang, 2009; Hayashi et al., 2009). Furthermore, Dally can bind Dpp and is proposed to regulate its extracellular distribution (Akiyama et al., 2008). We therefore visualised the localisation of endogenous Dally tagged with mCherry (Dally^mCh^). Like Dpp, we find an anteroposterior gradient of punctate extracellular Dally^mCh^ (Fig. 4B-Bi), suggesting that Dally binds Dpp and contributes to the formation of the Dpp gradient. However, this ligand distribution is incompatible with the classical view of Dpp signaling in the germarium, which predicts Dpp accumulation at the interface of GSCs and niche CpCs (Fig. S1B). We therefore addressed where Tkv activation occurs using a fluorescent reporter of ligand-receptor interaction, TIPF (Fig. 4C; Michel et al., 2011). TIPF is Tkv C-terminally tagged with YFP held in a non-fluorescent conformation, but upon ligand-receptor binding the YFP is released and adopts a fluorescent conformation. Using the endogenous fluorescence of germline expressed TIPF (*Bam27::TIPF*), we find the majority of active Tkv co-occurring with Dally^mCh^ (Fig. 4D-Di). This suggests that Dally-bound Dpp is a key source of ligand for GSCs and that signaling likely occurs on GSC projections that are extended into the niche.

To address whether Dally regulates Dpp distribution around the niche, the Gal80^ts^ system was used to temporally induce the knockdown or overexpression of Dally in niche cells. In adults raised at 18°C, Gal80^ts^ represses Gal4 activity and therefore expression of the associated transgene. Shifting adults to 29°C for 3 days causes Gal80^ts^ repression to enable transgene expression. When *dally^KD^* or *dally^OE^* adult flies are raised at 18°C, extracellular Dpp^mCh^ shows an anterior high gradient as seen in wildtype germaria (Fig. S5E-F). Inducing *dally* knockdown results in more equal levels of Dpp across the niche (Fig. 4E, Fig. S5E-Ei). Conversely, *dally* overexpression leads to greater anterior accumulation of Dpp^mCh^, resulting in a steeper gradient with a higher average anterior to posterior ratio (Fig. 4E, S5F-Fi). Together these results suggest that niche expressed Dally binds and sequesters Dpp away from the GSCs forming a reservoir of Dpp.

To provide further evidence that Dally-bound Dpp is the source of ligand for GSCs, we used the TIPF reporter to monitor Tkv activation following manipulation of *dally* expression in niche cells, as described above. Induction of *dally^KD^* expression at 29°C decreases the number of TIPF puncta compared to controls (18°C; Fig. 4F-Fi). Conversely, *dally* overexpression increases the number of niche TIPF puncta. These data are consistent with Dally-bound Dpp representing the major source of self-renewal signal for GSCs while the extension of projections likely enables access (Fig. 4G).

To directly determine if GSC projections allow access to the anterior reservoir of Dpp, we used live *ex vivo* imaging to monitor Dpp and Tkv localisation on GSC projections. An N-terminally tagged Tkv^YFP^ knock-in line (Lowe et al., 2014), while homozygous viable, was found to generate tumours of GSC-like cells (Fig. S5A-B), suggesting that Tkv regulation is impaired. Conversely, a C-terminally tagged Tkv^mCh^ knock-in line is homozygous viable and displays no germline phenotype (Fig. S5A-B) so was used hereafter. Firstly, the localisation of Tkv was monitored on APs using germline expression of *LifeAct.GFP* (Fig. 5A). Tkv^mCh^ is detected at the GSC-niche interface (Fig. 5A; yellow arrowhead) and around short APs (white arrowhead). Upon extension into the niche, Tkv^mCh^ localises along the projection (magenta arrowhead), suggesting that APs could function as Dpp signaling platforms. In addition, Tkv^mCh^ also decorates lateral GSC projections (Fig. S5G). Tkv^mCh^ was also visualised in *GFP.αTub* expressing germline stem cells (Fig. 5B). Here large puncta of Tkv^mCh^ were detected trafficking onto cytocensors and accumulating at the tip. The trafficking of larger Tkv^mCh^ puncta could be indicative of the active trafficking of Tkv^mCh^ onto cytocensors (e.g. by vesicular transport) in comparison with APs which may be more random. To determine whether these projections contact the Dpp reservoir, Dpp^mCh^ was visualised with *GFP.αTub* (Fig. 5C). Cytocensors form stable contacts with Dpp^mCh^ puncta. Finally, in order to determine whether Tkv activation occurs on GSC projections, we used germline expression of *UASp-FTractin.dTomato* to visualise actin filopodia alongside TIPF. In Figure 5D a single TIPF puncta is localised to the base of a pre-existing filopodium. The projection collapses and at 1:12 min has reformed. At this point a TIPF puncta appears at the tip of the AP and moves toward the base. GSC projections can therefore act as sites of Dpp signal transduction. Together these data are consistent with the GSCs dynamically localising Tkv onto cellular projections to permit access to Dpp.

**Figure 5.**
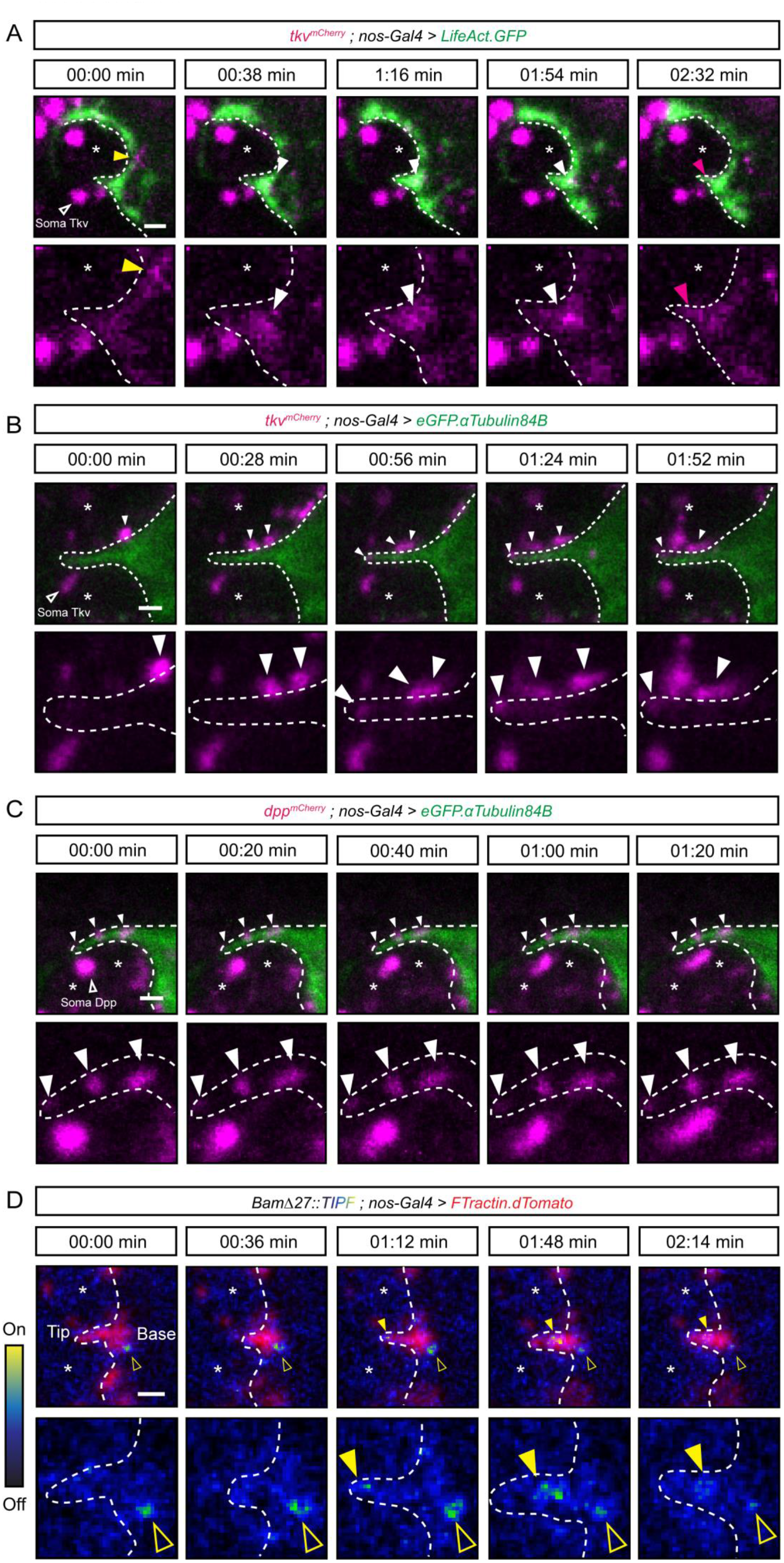
Tkv is trafficked onto GSC projections which act as signaling platforms. (A) Stills from a video of F-actin labelled with *LifeAct.GFP* and endogenous mCherry-tagged Tkv showing Tkv^mCh^ (yellow arrowhead) at the GSC-niche interface before accumulating at the base (white arrowhead) and then upon the AP (magenta arrowhead). (Bottom) shows closeup view of Tkv^mCh^ channel. (B) as in (A) showing the trafficking of Tkv^mCh^ puncta (white arrowheads) on a cytocensor labelled with *eGFP.αTub.* (Bottom) shows closeup view of Tkv^mCh^ channel. (C) as in (B) showing Dpp^mCh^ puncta (white arrowheads) statically associated with a cytocensor labelled with *eGFP.αTub.* (Bottom) shows closeup view of Dpp^mCh^ channel. (D) as in (A) showing active Tkv at the base of an FTractin-dTomato-labelled AP (TIPF fluorescence, open yellow arrowhead) and Tkv activation occurring on an AP (yellow arrowhead). (Bottom) shows closeup view of TIPF. TIPF is false-coloured as a heat-map for clarity. Projections are outlined by dashed lines. (*) CpCs. Scale bars = 1µm. See also Figure S5.

### Dia-regulates actin projection formation and Dpp signal reception

As Dpp signal transduction is observed on GSC projections, we next wanted to test their requirement for Dpp signaling. We firstly returned to the actin polymerising factor Dia which is necessary for GSC maintenance (Fig. 3). By raising *dia^KD^* flies at 18°C *shRNA* expression is repressed during larval development (Fig. 6A-B). Adults maintained at 18°C for 3 days do exhibit some GSC loss, suggesting there is leaky *shRNA* expression and highlighting the sensitivity of GSCs to *dia* levels. When adults are shifted to 25°C for 3 days, rapid GSC loss is observed (Fig. 6A-B), with many germaria containing single (Fig. 6A middle panel) or no GSCs (bottom panel). To determine if GSC loss is due to perturbed Dpp signaling, we visualised pMad. When *dia^KD^* expression is induced we do not observe a loss of pMad, instead neighbouring GSCs begin exhibiting greater variability in pMad levels. Typically, one GSC per niche experiences slightly higher levels of pMad than its neighbour (Fig. 6A top panel and C-D). This disparity may be due to uneven knockdown in neighbouring GSCs. Previous studies have shown that Dpp signaling mutant clones are outcompeted by neighbouring wildtype GSCs and replaced by symmetric cell division (Xie and Spradling, 1998). The disparity in pMad levels seen with *dia^KD^* expression could similarly be expected to promote such competition and lead to a ‘winner’ higher pMad GSC outcompeting the other. However, if this occurs here, we speculate that the block on cytokinesis due to low *dia* leads to single large polyploid ‘winner’ GSCs that occupy entire niches (Fig. 6E). We note that other functions of Dia could also contribute to GSC loss, such as polyploidy. We also rule out that GSC loss is due to reduced niche adhesion as Ecad levels at the GSC-niche interface increase following *dia* knockdown (Fig. 6F-G) consistent with disrupted endocytosis (Levayer et al., 2011, see below). Regardless of mechanism, we show that reduced *dia* expression leads to enhanced Dpp signaling in a subset of GSCs.

**Figure 6.**
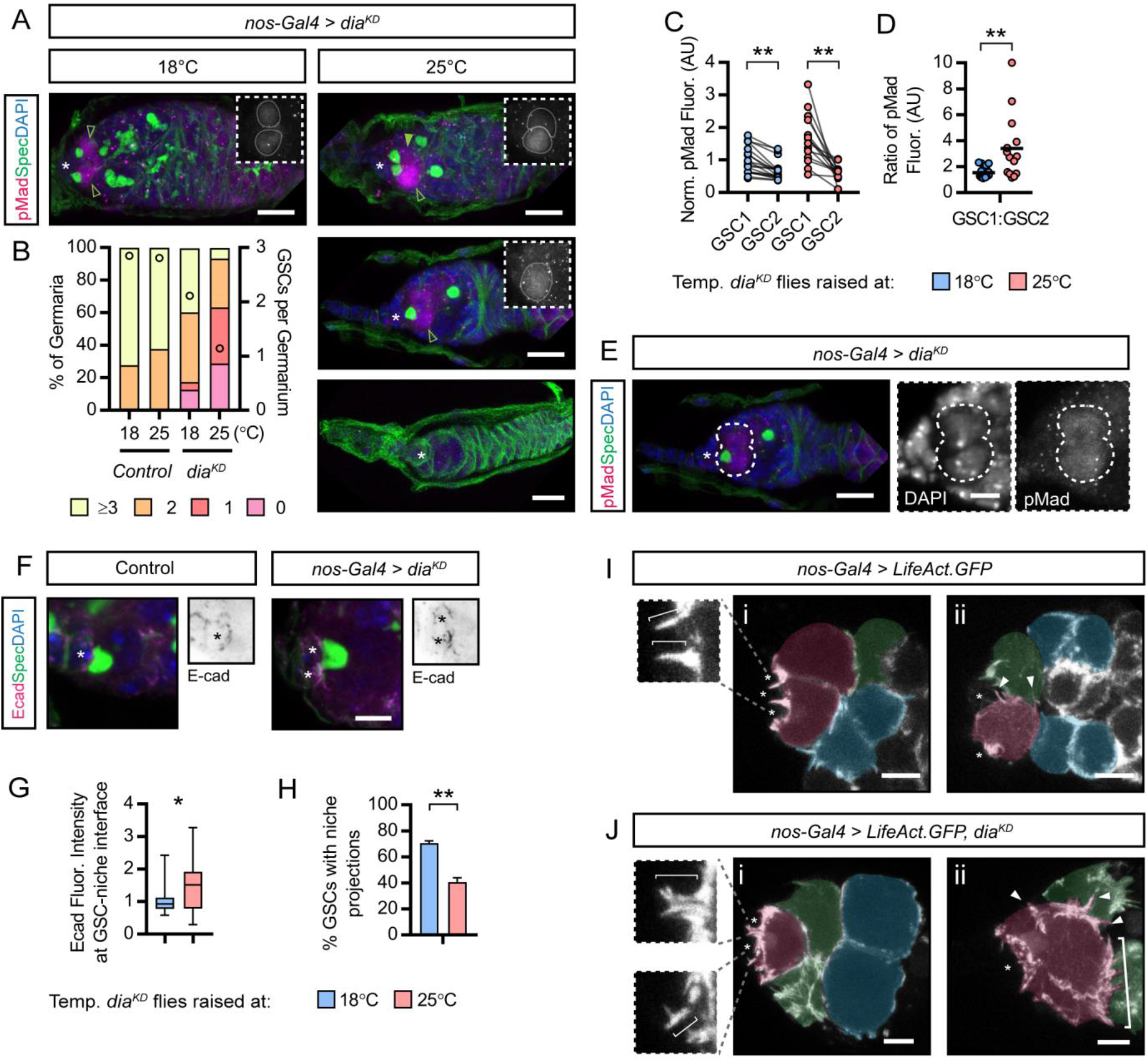
Knockdown of *dia* expression causes dysregulation of Dpp signaling and actin projections formation. (A) Germline-specific *dia^KD^* expression. Comparison of adults raised at 18°C for 3 days (inhibiting *shRNA* expression) or at 25°C for 3 days (inducing *shRNA* expression). (Insets) pMad staining reports the Dpp signaling response. Early germ cells are marked by the presence of the spectrosome (αSpectrin). (B) Histogram showing quantification of GSC numbers in (A) n=50 germaria for control, n≥142 for *dia^KD^*. (C) Comparison of pMad fluorescence shown in (A). GSC1 is the cell with the highest pMad and GSC2 has the lowest. n>15 germaria. (D) Ratio of pMad levels from (C). (E) A single polyploid *dia^KD^* expressing GSC that occupies an entire niche. (F) as in (A) showing Ecad in inverted black and white for clarity and (G) quantification in control (n=27) and *dia^KD^* (n=32). Median, 25^th^ and 75^th^ percentile and whiskers show minima and maxima. (H) Percentage of *dia^KD^* expressing GSCs that form actin-rich projections labelled with LifeAct.GFP. n>100 GSCs. (I-J) Stills showing actin projections labelled with LifeAct.GFP in wildtype (G) and *dia^KD^* expressing germ cells. Brackets in Iii label supernumerary projections and arrowheads label lateral projections. GSC (false coloured magenta), CB (green) and 2 or 4-cell cysts (blue). (*) CpCs. Scale bars = 5µm (A and F) or 2µm (G and H). *, p<0.05; **, p<0.001.

We next determined whether Dia regulates actin-projection formation using *LifeAct.GFP* co-expression. Upon *dia^KD^* expression the number of GSCs extending projections into the niche is decreased (Fig. 6H), whereas the projections that remain are abnormal. While wildtype projections are thin, finger-like filopodia (Fig. 6Ii), or broader lamellipodia (Fig. 2C), *dia^KD^-*GSCs extend branched, thick projections (Fig. 6Ji). Furthermore, their formation appears disorganised with the extension of supernumerary lateral projections (compare the APs (arrowheads) in Fig. 6Iii and the APs (arrowheads) and lamellipodial projections (bracket) in Fig. 6Jii). These results suggest that enhancing GSC projection formation could increase Dpp signal transduction.

### Cytoskeletal projections are necessary for Dpp signal activation and attenuation

If GSC projections are necessary for accessing the Dpp reservoir, inhibiting projection formation would be predicted to compromise Dpp signal reception, transduction and GSC fate. We first tested whether inhibiting projection formation with chemical inhibitors (Fig. 1I) disrupted receipt of Dpp in GSCs using the number of TIPF puncta around the niche as a readout. Germaria incubated *ex vivo* with DMSO maintained similar levels of signal activation after 30 and 90 mins (Fig. 7A-B). Incubation with CytoD, however, results in a significant reduction in TIPF puncta even after 30 mins (Fig. 7A-B) in agreement with a role for projections in Dpp signal reception. Incubation with nocodozole for 30 mins has no effect on the average number of TIPF puncta (Fig. 7A-B), suggesting that in the absence of cytocensors, GSCs are still able to access and receive Dpp through APs (Fig. 1I). However, there is a small but significant increase in the number of TIPF puncta following a 90 min nocodazole treatment (Fig. 7A-B). The overactive signaling observed in the absence of cytocensors raises the possibility that, in contrast to APs, they are necessary for signal attenuation.

**Figure 7.**
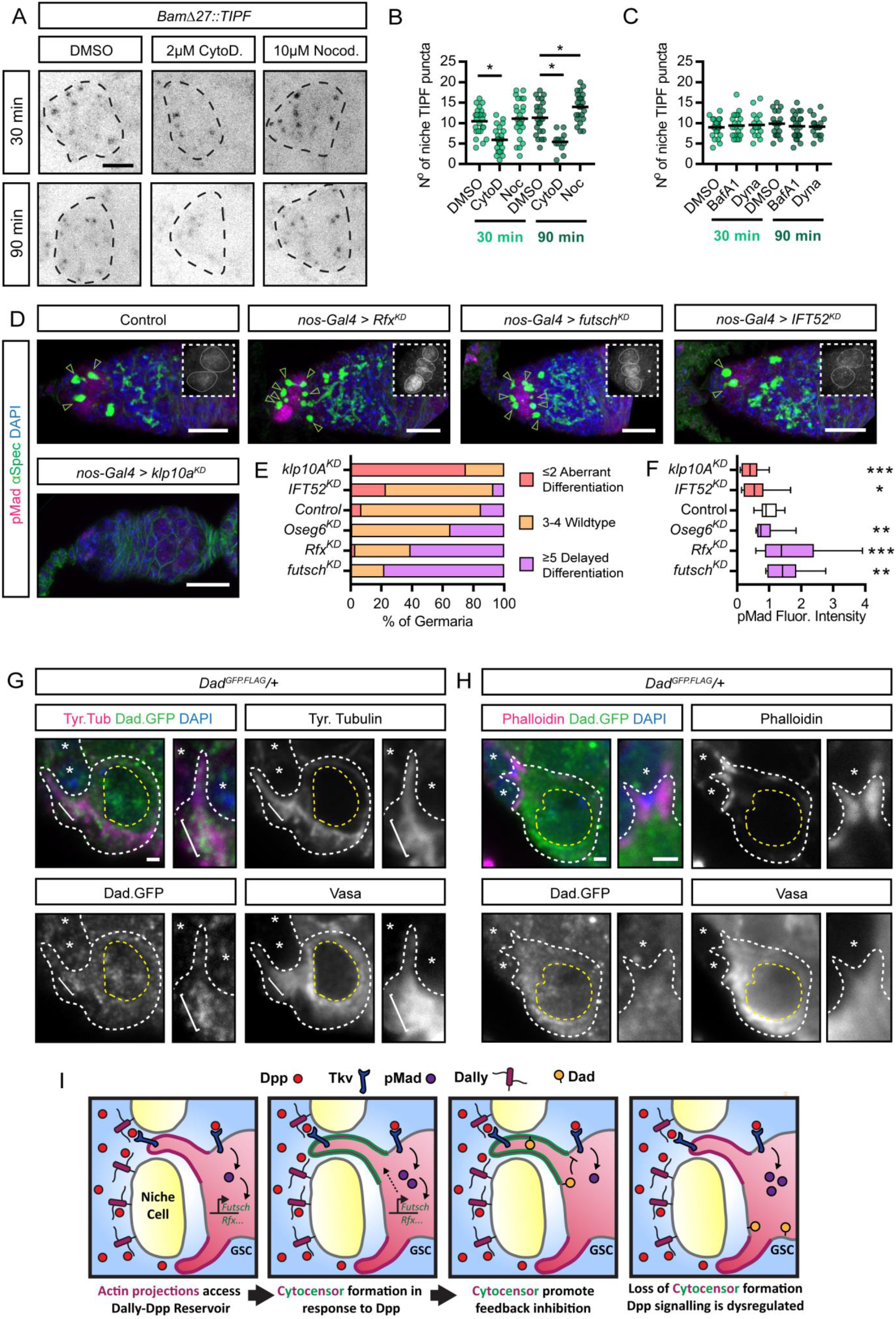
Cytocensors attenuate Dpp signaling to regulate GSC self-renewal and differentiation. (A) Endogenous TIPF fluorescence after 30 or 90 mins *ex vivo* drug treatment. (B-C) Scatter plots showing number of TIPF puncta per niche after treatment with (B) DMSO (n=24 each), 2µM CytoD (n=28 and 12, respectively), 10µM nocodazole (n=24 each) and (C) DMSO (n=24 and 19, respectively), 100nM BafA1 (n=22 and 23, respectively) and 100µM dynasore (n=19 each). (D) Germline-specific *shRNA e*xpression phenotypes. (Insets) pMad staining reports the Dpp signaling response. Early germ cells are marked by the presence of the spectrosome (open arrowhead) labelled by anti-αSpectrin. (E) Histogram showing quantification of GSC numbers in (B) and differentiation phenotype. n≥100 germaria. (F) Comparison of pMad fluorescence relative to controls in (D). Median, 25^th^ and 75^th^ percentile and whiskers show minima and maxima. n≥15 cells. (G-H) Immunofluorescence staining of a *Dad^GFP.FLAG^/+* germarium. White dashed line shows individual GSCs and yellow dashed line outlines the nucleus. Bracket indicates Dad.GFP concentrated at the base of a cytocensor. (I) Model illustrating the role of APs in promoting Dpp signal transduction and cytocensors in modulating levels through signal suppression. See text for details. Dashed lines in (A) outline the niche. Scale bar = 10µm (B) or 5µm (A), or 1µm (G). See also Figures S6 and S7. *, p<0.05; **, p<0.001; ***, p<0.0001. See also Figure S6 and S7.

An alternative interpretation of these data is that interfering with actin/MT polymerisation disrupts receptor endocytosis, trafficking, and/or degradation, all of which can influence BMP/Dpp signaling output (Ehrlich, 2016). To test this, we assessed the disruption to trafficking in the drug treatment timeframe and whether this would be sufficient to give the observed changes in TIPF puncta numbers (Fig. 7A-B). First we carried out a dextran uptake assay using fluorescently-tagged Dextran as a fluid-phase marker (Fig. S6A-E). While Dextran uptake is readily observed in controls (Fig. S6A), treatment with CytoD greatly reduces uptake (Fig. S6B), consistent with observations that inhibiting actin polymerisation blocks endocytosis (Mortensen and Larsson, 2003). A similar result is obtained following treatment with Dynasore (Fig. S6C), a potent inhibitor of dynamin-dependent endocytosis (Macia et al., 2006). Following nocodazole treatment we observe increased dextran accumulation in smaller vesicles (Fig. S6D), consistent with the inhibition of MT polymerisation disrupting endosomal trafficking/maturation (Bayer et al., 1998). A similar result is observed upon treatment with bafilomycin A1 (BafA1) (Fig. S6E), an inhibitor of endosomal/lysosomal maturation.

As these data show that incubating ovaries with CytoD and nocodazole perturbs endocytosis and trafficking, we next determined whether these additional effects of disrupted actin/MT polymerisation could account for the altered Tkv activation (Fig. 7A-B). To this end, we used Dynasore and BafA1 to specifically assay the effect of disrupting endocytosis and degradation on the number of niche TIPF puncta. In both cases, we observe no effect after 30 or 90 mins of treatment (Fig. 7C and S6F). These data suggest that the reduced/enhanced Tkv activation observed in Fig. 7A-B is not due to reduced Tkv endocytosis or trafficking/degradation.

To complement the above data, we specifically investigated the effect of CytoD and nocodazole treatment on the endocytosis of Tkv^mCh^ by monitoring its co-localisation with Rab5^YFP^, as early endosomal trafficking of Tkv in Rab5^+^ endosomes is known to enhance Dpp signal response (Gui et al., 2016). Only a small proportion of the Rab5^+^ endosomes per GSC are also Tkv^mCh^ positive in controls (Fig. S6G-I). Incubation with CytoD significantly decreases the number of Rab5^+^ endosomes, indicative of inhibited endocytosis (Fig. S6G-H; Mortensen and Larsson, 2003). However, this has little effect on the average number of Tkv^mCh^-positive Rab5^YFP^ puncta per GSC (Fig. S6I) compared to the control. Incubation with nocodazole significantly increased the number of Rab5^+^ endosomes (Fig. S6H), however, this also had little effect on the average number of Tkv^mCh^-positive Rab5^YFP^ puncta per GSC (Fig. S6I). These data provide further evidence that the effect of CytoD or nocodazole on early endosomal trafficking of Tkv is insufficient to explain the altered Tkv activation observed.

We also addressed whether inhibition of actin polymerisation leads to the loss of Ecad-based AJs which are necessary for GSC maintenance (Song et al., 2002). Following 90 mins of CytoD or nocodazole treatment GSCs maintain contact with the niche CpCs (Fig. S6J). Additionally, the spectrosome remains anteriorly anchored to the niche interface suggesting that polarity is maintained (Fig. S6J; Deng and Lin, 1997). Nevertheless, we tested the effect of disrupting Ca^2^-dependent cadherin binding by incubating germaria *ex vivo* with the Ca^2+^ chelator EGTA. A 90 min treatment with EGTA induces germline Ecad internalisation (Fig. S6K). However, EGTA treatment has no effect on the number of TIPF puncta per niche after 90 and even 150 mins (Fig. S6L-M). This shows that reduced niche adhesion is not sufficient to reduce Dpp signal transduction. We therefore favour the conclusion that reduced TIPF activation following CytoD treatment is due to loss of projections rather than its effect on Tkv trafficking or GSC adhesion.

Next, we further investigated cytocensor function through genetic manipulation by disrupting their formation through the germline-specific knockdown of factors described in Fig. 3. pMad staining and the number of early germ cells were used to assay effects on Dpp signal transduction and GSC maintenance/differentiation, respectively. RNAi-knockdown of *IFT52* or *klp10A*, which increases the frequency of projection formation or enhances the growth of cytocensors, respectively, leads to an increased rate of GSC loss (Fig. 7D-E) and decrease in pMad levels (Fig. 7F). This suggests GSCs aberrantly differentiate due to reduced Dpp signalling. We also observe a decrease in Ecad levels at the GSC-niche interface in *klp10^KD^* GSCs (Fig. S7A-B). However, it is not possible to determine whether this is a differentiation-associated loss (Shen et al., 2009) or due to a cytocensor-independent function of Klp10A in Ecad localisation, although we favour the former as we observe continued Dpp signalling following loss of adherens junctions by EGTA treatment (Fig. S6L-M). We speculate that enhancing cytocensor formation or length is detrimental to the maintenance of pMad activation, such as through mis-trafficking of Tkv on long projections. However, as kinesin-like proteins have also been shown to shuttle endosomes, we cannot exclude that defective endosomal trafficking contributes to the lower pMad level in GSCs and their loss.

A weak delay in differentiation, typified by accumulation of ectopic early germ cells, is observed upon expression of *Oseg6^KD^* (Fig. 7E). This is stronger upon knockdown of *futsch* or *Rfx* (Fig. 7D-E), which both disrupt cytocensor formation (Fig. 3). Consistent with this delayed differentiation, in both cases we detect a significant increase in GSC pMad levels (Fig. 7F). These data show that when cytocensor formation is disrupted by knockdown of specific genes or chemical inhibition, Dpp reception is enhanced suggesting that cytocensors have an additional function, which is to attenuate Dpp signaling.

An alternative interpretation of the above data is that genetic knockdown of the regulators leads to a prolonged disruption of endocytosis, which alters GSC/CB fates through impaired BMP signal activation in GSCs and/or altered E-cadherin accumulation at the GSC-niche interface. We first addressed this by measuring dextran uptake following knockdown of cytocensor-regulatory factors. Knockdown of *dia*, a known regulator of endocytosis (Levayer et al., 2011), reduces dextran uptake (Fig. S7C-D). However, no reduction or gross effect on dextran uptake is observed upon knockdown of the cytocensor regulators. Nonetheless, we directly addressed the effect of genetically inhibiting endocytosis on GSC/CB fates. We used germline knockdown of *shibire* (*shi*), encoding Dynamin, to inhibit dynamin-dependent endocytosis (Fig. S7E-F). This results in delayed differentiation of cysts and accumulation of cysts and developing nurse cells (yellow dashed line) within germaria (Fig. S7E). *shi^KD^* efficiently inhibits dextran uptake in GSCs (Fig. S7C-D) and results in increased Ecad levels at the GSC-niche interface (Fig. S7G-H), consistent with reduced Ecad endocytosis, but no change in pMad levels (Fig. S7E-F). Together these data suggest that the Dpp signal response in GSCs is independent of receptor endocytosis, while conversely GSC-niche adhesion and germ cell differentiation are regulated by dynamin-dependent endocytosis. As increased Ecad does not alter GSC number (Pan et al., 2007) and the phenotypes associated with knockdown of cytocensor regulators in Fig. 7 are associated with alterations in pMad levels, we favour the interpretation that the phenotypes are due to impaired reception or attenuation of the Dpp signal, rather than secondary effects on endocytosis.

We also investigated whether Rfx and Futsch could regulate the BMP signaling response between GSCs and their daughters independently of cytocensors. GSCs undergo an unusual cell cycle with delayed cytokinesis such that the GSC remains attached to its daughter pre-cystoblast (pCB) until G2 phase. Tkv activity is terminated in the pCB through the asymmetric upregulation of the Fused kinase, which is essential for differentiation by targeting Tkv for degradation (Xia et al., 2010, 2012). This results in the generation of a pMad gradient between the GSC-pCB with 20-30% of pCBs displaying half the level of pMad observed in the GSC, whereas the majority of pCBs rapidly reduce pMad levels to less than a quarter (pMad^-^) that of the GSC to enable differentiation (Fig. S7I-K; Xia et al., 2012). Upon knockdown of *futsch* or *Rfx*, we observe more pMad^+^ pCBs (Fig. S7I-K), however the pMad gradient (relative pMad ratio between GSC and pCB) remains the same as observed for pMad^+^ pCBs in control germaria. This suggests that following *futsch* or *Rfx* knockdown, the circuitry regulating Dpp signal termination in pCBs remains functional while the GSC exhibits enhanced Dpp signal reception. We therefore propose that the regulation of the Dpp signal response in GSCs by Futsch and Rfx is more likely through the promotion of cytocensor formation, which attenuates signaling levels.

Finally, to address how Dpp signaling may be attenuated we tested the hypothesis that the cytocensor provides a compartment for the coordinated concentration of both signaling machinery and antagonists. The classic Dpp target gene and inhibitory Smad, Dad, is expressed in GSCs and reduces Dpp signaling levels (Casanueva and Ferguson, 2004; Xie and Spradling, 1998). Immunofluorescence staining of transgenic GFP-tagged Dad shows that it is diffuse throughout the cytoplasm and nucleus and concentrates at the base of cytocensors (Fig. 7G) but not APs (Fig. 7H), consistent with a cytocensor-specific signal attenuating property. In conclusion, we propose that the role of GSC projections is two-fold; projections allow the receipt of secreted Dpp held away from the GSCs within a niche reservoir, while cytocensors enable the concerted localisation of signaling machinery and antagonist/s to modulate signaling levels (Fig. 7I).

## DISCUSSION

Here we present data describing the changing transcriptome of GSCs as they transition from self-renewal to differentiation. Genes up-regulated in GSCs are associated with gonad development, chromatin organisation and transcriptional regulation. This is consistent with germline differentiation being accompanied by a reduction in transcriptional activity and altered chromatin organisation (Flora et al., 2018; Zhang et al., 2014). Genes upregulated in CBs include those encoding factors involved in cellular metabolism, growth and protein production. This can be rationalised with CB biology as, upon exiting the stem cell niche, the CB quickly undergoes 4-rounds of mitosis to generate a 16-cell cyst and this transition is associated with an increase in general and mitochondrial protein synthesis (Sanchez et al., 2016; Teixeira et al., 2015). We also identified Dpp target genes in GSCs and have elucidated the function of two positive Dpp targets, *Rfx* and *futsch*, which promote synthesis of cytocensors along with other MT-associated genes in the GSC transcriptome.

Based on our data, we propose a model whereby GSCs synthesise APs that access a Dally-bound reservoir of Dpp (Fig. 7I). In support of this, short-term inhibition of AP formation significantly reduces active Tkv levels, whereas increasing AP branching and disorganisation (*dia^KD^*) is associated with increased pMad levels. We propose that, following activation of Dpp target genes, such as *Rfx* and *futsch*, these APs develop into cytocensors (Fig. 7I) and provide evidence that, like the APs, these cytocensors access Dpp localised away from the GSCs.

GSC projections act at much shorter range than that typically associated with signaling filopodia, which in many contexts are associated with the regulation of long-range signal transduction. For example, cytonemes in the larval wing disc and dorsal air sac primordium drive long-range Dpp signaling (Wilcockson et al., 2017). The advantage of the unusual GSC-niche architecture may be two-fold. Firstly, it concentrates the potent self-renewal signal further away from the GSCs, guarding against ectopic Dpp diffusion that would disrupt GSC differentiation. Secondly, and we suggest more importantly, our data provide evidence that the cytocensors allow GSCs to actively regulate their Dpp signaling levels by both collecting Dpp and attenuating signal transduction. Genetic or chemical perturbation of cytocensors leads to increased active Tkv and pMad (Fig. 7I), suggesting that these cytocensors promote feedback inhibition to maintain signal transduction at a threshold that facilitates differentiation following GSC division. Mathematical modelling has previously shown that the pMad concentration prior to division is important so that levels fall below a critical threshold in the GSC daughter, allowing *bam* derepression and differentiation (Harris et al., 2011).

We speculate that cytocensors facilitate Dpp signal termination by acting as a hub where GSCs can concentrate signaling machinery and antagonists, such as Dad, to efficiently modulate pMad levels. Similarly, the Dad homolog Smad7 has been shown to localise to the base of primary cilia where its proposed to inhibit TGFβR-Smad interaction to modulate signaling levels (Pedersen et al., 2016). This signal attenuating activity of female cytocensors is in stark contrast to the role of male MT-nanotubes, which increase the stem cell-niche interface area to promote Dpp signal transduction. Perturbing nanotube formation decreases pMad levels, increasing the rate of competition-induced loss. Furthermore, male and female projections exhibit several structural differences. Male projections are MT-based static structures formed independently of actin, and most GSCs (∼80%) extend one or more of these projections into the body of neighbouring niche cells (Inaba et al., 2015). Female cytocensors, on the other hand, are rich in both MTs and actin, are relatively dynamic, transient structures, and are much less frequently found (∼40% of GSCs). However, we propose that all GSCs will synthesise cytocensors, but at different times as they form in response to increasing Dpp signalling levels as part of a feedback mechanism. The distinct nature of male and female projections may be due to differing requirements for Dpp signaling in stem cell maintenance or more limiting Dpp levels and the competition between GSCs and somatic cyst stem cells for niche occupancy in the testis (Greenspan et al., 2015).

Given our evidence for signal transduction through GSC projections, this raises the question as to whether GSCs can receive Dpp in their absence. Monitoring Dpp-Tkv interaction following CytoD treatment shows that TIPF puncta are still present up to 90 min after treatment. This could be due to incomplete loss of APs or may suggest that diffusing Dpp drives low-level signal transduction. Alternatively, the maintenance of a few puncta could be due to perdurance of the TIPF reporter or disrupted trafficking/endocytosis of Tkv that was active prior to treatment (Lamaze et al., 1997). The identification of additional actin regulators involved in this process will be necessary to determine the absolute requirement for APs in GSC Dpp signaling. One putative factor is the small GTPase Rac1, a key regulator of the formation and identity of APs, which concentrates at the GSC-niche interface (Lu et al., 2012). Rac1 is necessary for long-term GSC maintenance and was suggested to promote BMP signal transduction.

We also find lateral GSC and CB projections decorated with Tkv^mCh^. It has been shown that differentiating germ cells remain highly sensitive to ‘leaky’ Dpp, while ectopic Dpp can induce germ cell dedifferentiation (Xie and Spradling, 1998). The formation of these projections may therefore promote the sensitisation of germ cells to Dpp signaling, enabling their dedifferentiation to replace GSCs lost during aging or stress (Liu et al., 2015).

Dynamic signaling projections enables the receipt or delivery of signaling molecules over large distances or between cells of different tissues. The ability of signaling projections to modulate signal transduction may be of particular importance to adult stem cells. These are cells that need to be sensitive to local and systemic signaling while still maintaining their own plasticity. While signaling projections are common during development, it remains to be determined whether they are also frequently found in adult stem cells. However, in the murine gut, intestinal stem cells extend apical processes that reach in between neighbouring Paneth cells that constitute a key part of the intestinal stem cell niche (Barker et al., 2007) and Lgr4/5 has been shown drive formation of cytoneme-like projections *in vitro* (Snyder et al., 2015). It will be interesting to determine the role of these projections in stem cell signaling and whether cytocensors are found in other developmental contexts.

## ACKNOWLEDGEMENTS

We thank Ryo Hatori, Thomas Kornberg, Georgios Pyrowolakis, the Bloomington Drosophila Stock Center and the Developmental Studies Hybridoma Bank for flies and/or antibodies. We thank Catherine Sutcliffe and Rosalind Wilkes for technical assistance, University of Manchester Genomic Technologies Facility and Ping Wang for analysis of the RNA-seq data, Jens Januschke for advice on live imaging and Joseph Morgan for helpful discussions. This work was supported by a BBSRC PhD Studentship.

## AUTHOR CONTRIBUTIONS

Conceptualisation, S.G.W, H.L.A; Methodology, S.G.W, H.L.A; Investigation, S.G.W; Formal Analysis, S.G.W; Writing, S.G.W, H.L.A; Supervision, H.L.A; Funding Acquisition, H.L.A.

## METHODS

### Experimental Model and Subject Details

Flies were grown at 25°C using standard procedures. The following fly lines were used in this study; *vasa.eGFP.HA* and *tkv^eYFP^* (Kyoto Stock Center), *bam.GFP* (Chen and McKearin, 2003), *UASp-tkv^QD^* (Tanimoto et al., 2000), *dally^mCh^* (Norman et al., 2016), *BamΔ27::TIPF* (Michel et al., 2011), *dpp^HA^* (Shimmi et al., 2005), and CRISPR knock-in *dpp^mCh^* (Fereres et al., 2018) and *tkv^mCh^* (gift from T. Kornberg). *dpp^mCh^* was generated by tagging Dpp after amino acid 465, as previously described by Entchev et al. (2000), and *tkv^mCh^* was generated by replacing the STOP codon of endogenous *tkv* with the *mCherry* coding sequence. Both lines are homozygous viable, fertile and display no germline phenotype. The following were obtained from Bloomington Stock Center; *dpp^hr92^*, *tkv^7^, mad*^1-2^, *med^13^*, *dad^GFP.FLAG^, UAS-dpp^GFP^, bab1-Gal4, tub-Gal80^ts^, nos-Gal4::VP16*, *UASp-eGFP.αTubulin84B*, *UASp- Actin42A.eGFP*, *UASp-Actin5c.mRFP*, *UASp-LifeAct.eGFP*, *UASp-FTractin.dTomato* and *UASp-shRNA* lines. Germline-specific RNAi was carried out by crossing *UASp-shRNA* fly lines to those carrying the germline driver *nos-Gal4::VP16.* For RNAi, flies were moved to 29°C for 1 week post-eclosion before dissection to enhance the knockdown. For temporal control of *dia^KD^*, *shi^KD^* or *Gal80^ts^* flies were raised at 18°C and then either kept at 18°C or moved to 25°C or 29°C for 3 days before dissection. For all drug/EGTA treatments, 3-5 day old flies were used for all samples.

### RNA-seq

For each RNA sample 300-400 ovary pairs were dissected from 3-5 day old flies. Ovaries were dissected into 1x PBS on ice and incubated in 5 mg/ml collagenase IV in PBS (Worthington Biochemicals) at RT for 45 min to dissociate the tissue. Cells were washed in PBS and filtered through a 40 μm nylon mesh to remove debris before fluorescence-activated cell sorting (FACS) based on the expression of *vasa.GFP* or *bam.GFP* using a FACSAria™ Fusion cell sorter (Diva 8 Software; BD Biosciences). Cells were sorted into 1x PBS on ice and RNA was isolated using TRIzol (Invitrogen). Total RNA was processed and sequenced by the Genomic Technologies Core Facility (University of Manchester) on the Illumina Genome Analyser II. Reads were mapped to the *Drosophila* genome (dm3) and gene counts were analysed using HTSeq. DESeq2 was used to calculate differential expression between genotypes and differentially expressed genes were determined to be log_2_ fold change >0.5 more highly expressed than the other genotype with a p<0.05. GO term analysis was carried out using the Gene Ontology Consortium with Fisher’s Exact with FDR multiple test correction. RNA-seq data are available from ArrayExpress with the accession number E-MTAB-7063.

### *ex vivo* live imaging

For live imaging 3-5 day old flies were dissected in Schneider’s Insect Media (Thermofisher) supplemented with 10% Fetal Bovine Serum (FBS) (Thermofisher), 1% (w/v) penicillin/streptavidin (Sigma) and individual ovarioles were separated and the overlying muscle removed to reduce movement during imaging. Ovarioles were mounted on a 35mm glass bottom tissue culture dish (World Precision Instruments) in a drop of Schneider’s Insect Media (10% FBS, 1% pen/strep) supplemented with 10 mg/ml fibrinogen (Millipore) which is spread across the glass bottom well before adding 1ul of thrombin (10 U/ml; GE Healthcare Lifesciences) to clot the fibrinogen. Schneider’s Insect Media (10% FBS, 1% pen/strep) supplemented with 200 mg/ml Human insulin (Sigma) was added and ovarioles were maintained at room temperature for 2-3 hours.

Germaria were imaged using a Leica TCS SP8 AOBS inverted microscope using a HC PL APO CS2 motCORR 63x/1.2 water objective with 5-6x confocal zoom with a pinhole size of 1 AU, scan speed 400 Hz unidirectional, format 512 × 512, 2x line averaging, with 10-20 *z*-stacks taken at 0.75 μm intervals every 30-60 seconds. Imaging was carried out at room temperature for 1-3 hours and images were subsequently analysed using Fiji. For later analysis, a brightfield image was taken to localise the GSCs and niche.

### Immunofluorescence imaging

Ovaries were fixed in 4% formaldehyde in PBT (1x PBS, 0.1% Triton X-100) for 15 minutes, washed three times for 15 minutes in PBT and blocked in 10% Bovine serum albumin (BSA) in PBT for 30 minutes before overnight incubation with primary antibodies in 10% BSA in PBT at 4°C. Ovaries were then washed four times in PBT over an hour and incubated with secondary antibodies for 2 hours at room temperature. They were then washed twice for 15 minutes in PBT and once for 15 minutes in PBS before mounting in Prolong Gold Antifade with DAPI (Invitrogen). For phalloidin staining, ovaries were incubated for 30 minutes (during the second PBT wash before mounting) and washed twice for 15 minutes with PBT before mounting. Fixed germaria were imaged using a Leica TCS SP5 AOBS inverted microscope using an HCX PL APO 63x/1.4 oil objective or PL APO 100x/1.4 oil objective.

For visualising MT projections, flies were fixed in PEM buffer (80 mM PIPES (Sigma), 1 mM MgCl_2_ (Sigma), 5 mM EGTA (Sigma), pH 7.4) with 4% formaldehyde and 2 μM paclitaxel (Sigma) for 30 minutes and rinsed twice in PEM buffer and washed in 3 times for 15 minutes in PBT (1x PBS, 0.1% Triton X-100) before blocking in 10% BSA in PBT for 30 minutes. Protocol was followed as outlined above.

For visualising extracellular proteins only, ovaries were fixed in PEM buffer with 4% formaldehyde for 15 minutes, washed in PEM three times for 15 minutes and blocked for 30 mins in 5% BSA in PEM. Ovaries were incubated with primary antibodies in 5% BSA in PEM for 3 hours at room temperature, washed four times in PEM over an hour before incubation with secondary antibodies for 2 hours at room temperature. The ovaries were washed three times in PEM and mounted in Prolong Gold Antifade with DAPI. Antibodies were used at higher concentrations; Rt anti-DCAD2 (DSHB, 1:10), Rb anti-mCherry (ab183628, 1:50), Ms anti-HA 12CA5 (Roche, 1:50) and Rb anti-GFP (ab6556, 1:50).

To visualise endogenous TIPF and mCherry fluorescence, ovaries were fixed in PEM buffer with 4% formaldehyde for 15 minutes, before washing with PEM three times for 15 minutes. Ovaries were mounted in Prolong Gold Antifade and immediately imaged.

Antibodies used include; Rb anti-GFP (ab6556, 1:250), Goat anti-GFP (ab6673, 1:500), Rb anti-mCherry (ab183628, 1:500), Ms anti-RFP (ab65856, 1:250), Ms anti-αSpectrin 3A9 (DSHB, 1:50), Rt anti-DCAD2 (DSHB, 1:50), Ms anti-Futsch 22C10 (DSHB, 1:50) Rb anti-pSmad3 (ab52903, 1:500), Rt anti-αTubulin [YL1/2] (ab6160, 1:200), Ms anti-acetylated αTubulin [6-11b-1 (ab24610, 1:500), anti-γTubulin (Sigma GTU-88, 1:250), Rb anti-Vasa (Santa Cruz, 1:500). Secondary antibodies and other stains used were Alexa Fluor 488 Donkey anti-Rb, Alexa Fluor 488 Goat anti-Chk, Alexa Fluor 555 Donkey anti-Ms, Alexa Fluor 594, Donkey anti-Rt, Alexa Fluor 647 Donkey anti-Rt, Alexa Fluor 647 Donkey anti-Rb, Alexa Fluor 633-conjugated wheat germ agglutinin and Alex Fluor 488-conjugated Phalloidin (Thermofisher).

### Pharmacological inhibition

For *ex vivo* treatment with actin and MT depolymerising drugs, 5 ovary pairs per treatment were dissected in PBS and collected in Schneider’s Insect Media (10% FBS, 1% pen/strep). Ovaries were incubated with either the vehicle DMSO (Sigma), 2µM CytoD (Sigma), 10µM nocodazole (Sigma), 100µM dynasore (Sigma) or 100nm bafilomycin A1 (Sigma) for 30 mins or 90 mins at 25°C. Ovaries were rinsed with PBS and fixed. Immunostaining protocols were followed as previously described.

For EGTA treatment, 5 ovary pairs (*vasa.GFP* expressing females) per treatment were dissected in PBS and collected in Schneider’s Insect Media without FBS (1% pen/strep). Ovaries were treated with either H_2_O or 6mM EGTA for 90 or 150 mins. Ovaries were rinsed with PBS and fixed. Extracellular immunostaining protocol was followed as previously described. Endogenous Vasa.GFP expression was used for orientation when imaging.

Dextran uptake assays were performed by incubating ovaries (*shRNA* expressing or *vasa.GFP* expressing) with drugs in 100μl of media as described above for 30 mins prior to adding 3kDa Dextran-Texas Red or 10kDa Dextran-Alexa Fluor 488 (2 mg/ml; ThermoFisher) and incubating for further 60 mins. Ovaries were rinsed three times in PBS and fixed in 4% formaldehyde in PBS for 15 mins before mounting in Prolong Gold Antifade. For *shRNA* expressing germaria ovaries were incubated with Alexa Fluor 633-conjugated wheat germ agglutinin (5 μg/ml) for 15 mins before washing and mounting in Prolong Gold Antifade.

### Quantification of projection length, thickness and dynamics

Cytocensor length was measured using the line tool in Fiji to measure from the tip of the projection to the base. Thickness was measured at the base. For actin projections dynamics, live images were maximum projected and analysed in Fiji. Length was measured using the line tool to measure from the tip of the projection to the base at the point at which it achieves its maximum length. Extension speed was measured as the time taken from the point of projection nucleation and reaching maximum length. Retraction speed is defined as the time it takes a projection to completely collapse after reaching its maximum length. Lifetime is measured as the total amount of time the projection is visible for. The two-dimensional angle of nucleation was measured using the angle tool in Fiji to draw a line from the centre of the niche, determined from a brightfield view of the germarium, to the centre of the stem cell or cystoblast.

### Quantification of BMP signaling response and germ cell number

To measure pMad fluorescence, z-stacks (sum of slices) were generated at 0.75µm intervals of all 10-12 slices incorporating individual GSCs. Using the draw tool in Fiji, a circle encompassing the GSC was drawn and the integrated fluorescence density (IFD) was taken and background subtracted (IFD – (average mean intensity of background × area of the region sampled)). All results were then normalised to controls. Quantification of germ cell numbers was carried out using spectrosome staining. GSCs were classed as germ cells localised at the anterior tip of the germarium in contact with the niche and containing an anteriorly anchored spectrosome. Early germ cell quantification included GSCs and any additional round spectrosome containing cells. To count TIPF puncta, a brightfield image of the germarium was used to locate the niche and TIPF puncta were manually counted on Fiji. To measure the relative ratio of pMad between GSC-pCB pairs, z-stacks (sum of slices) were generated at 0.75µm intervals of all 10-12 slices incorporating individual GSCs. Using the draw tool in Fiji, a circle encompassing the GSC was drawn and the IFD was used to determine the ratio of pMad fluorescence. Quantification of Rab5+ and Tkv+ endosomes in GSCs was done manually.

### Quantification of Ecad levels

To measure Ecad fluorescence, z-stacks (sum of slices) were generated at 0.75µm intervals of all 8-12 slices incorporating individual GSCs-niche contact point. Using the draw tool in Fiji, a line was drawn over the area of contact between individual GSCs and neighbouring CpCs using α-Spectin staining which outlines the CpCs. Fluorescence intensity was calculated the same as above using nuclear intensity as background. All results were then normalised to controls. n indicated individual GSC-niche contacts from n>10 germaria from at least 2 biological replicates.

### Quantification of dextran uptake assay

To determine the size of dextran^+^ vesicles, 5 *z-*slices taken of entire germaria at 1µm intervals was used and maximum projected. The resulting images were background subtracted and thresholded to generate a binary image using Fiji. The area of each dextran^+^ vesicle within the region of the germarium, including both germline and somatic cells, was then measured using analyse particles setting a minimum size of 0.04µm^2^. The number of dextran puncta per GSC was manually counted as the total number of puncta within the anterior GSC region, outlined by wheat germ agglutinin, and divided by the total number of GSCs present (2-3).

### Statistical Analysis

Statistical comparisons were performed using two-tailed Student’s t-tests, one-way ANOVA with multiple comparisons or paired t-test (Fig. 6C) using GraphPad Prism and Microsoft Excel. Statistical significance was assumed by p<0.05. Individual p-values are indicated. Data are represented by the mean and standard deviation unless otherwise stated.

## SUPPLEMENTAL INFORMATION

**Figure S1.**
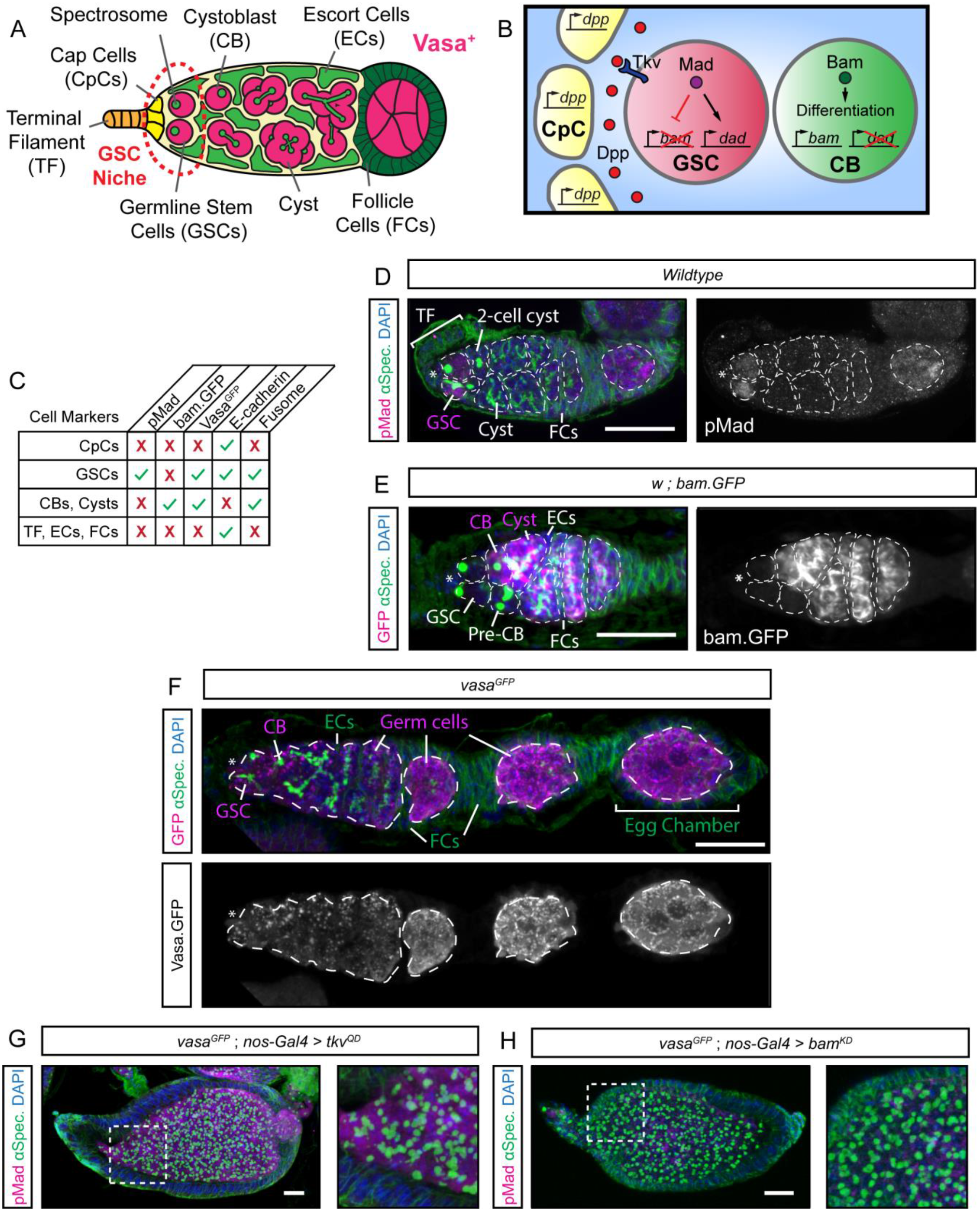
The *Drosophila* ovarian germline. Refers to Figure 1. (A) Cartoon depicting the structure of the germarium and GSC niche. The germline is indicated by Vasa expression (magenta). Niche cells are outlined by the dashed line. (B) Cartoon depicting the regulation of GSC self-renewal and differentiation by niche Dpp signalling. (C) Table indicating cell marker expression in all the cells of the germarium. (D) GSCs are identifiable as anteriorly localised pMad^+^ germ cells containing a single, round spectrosome. (E) Differentiating germ cells and cysts are identifiable by the expression of *bam* (shown here by a *bam.GFP* reporter). (F) All germ cells are marked by the expression of *vasa* (here a *vasa.GFP* reporter). (G) Germline-specific expression of constitutively active Tkv (Tkv^QD^) generates tumours of pMad^+^ GSC-like cells with single, round spectrosomes. (Inset) closeup view of boxed region. (H) Germline-specific expression of *bam^KD^* generates tumours of pMad^-^ GSC-like cells with single, round spectrosomes. (Inset) closeup view of boxed region. Scale bar = 5µm. CpCs (*). Dashed lines mark individual germ cells and cysts.

**Figure S2.**
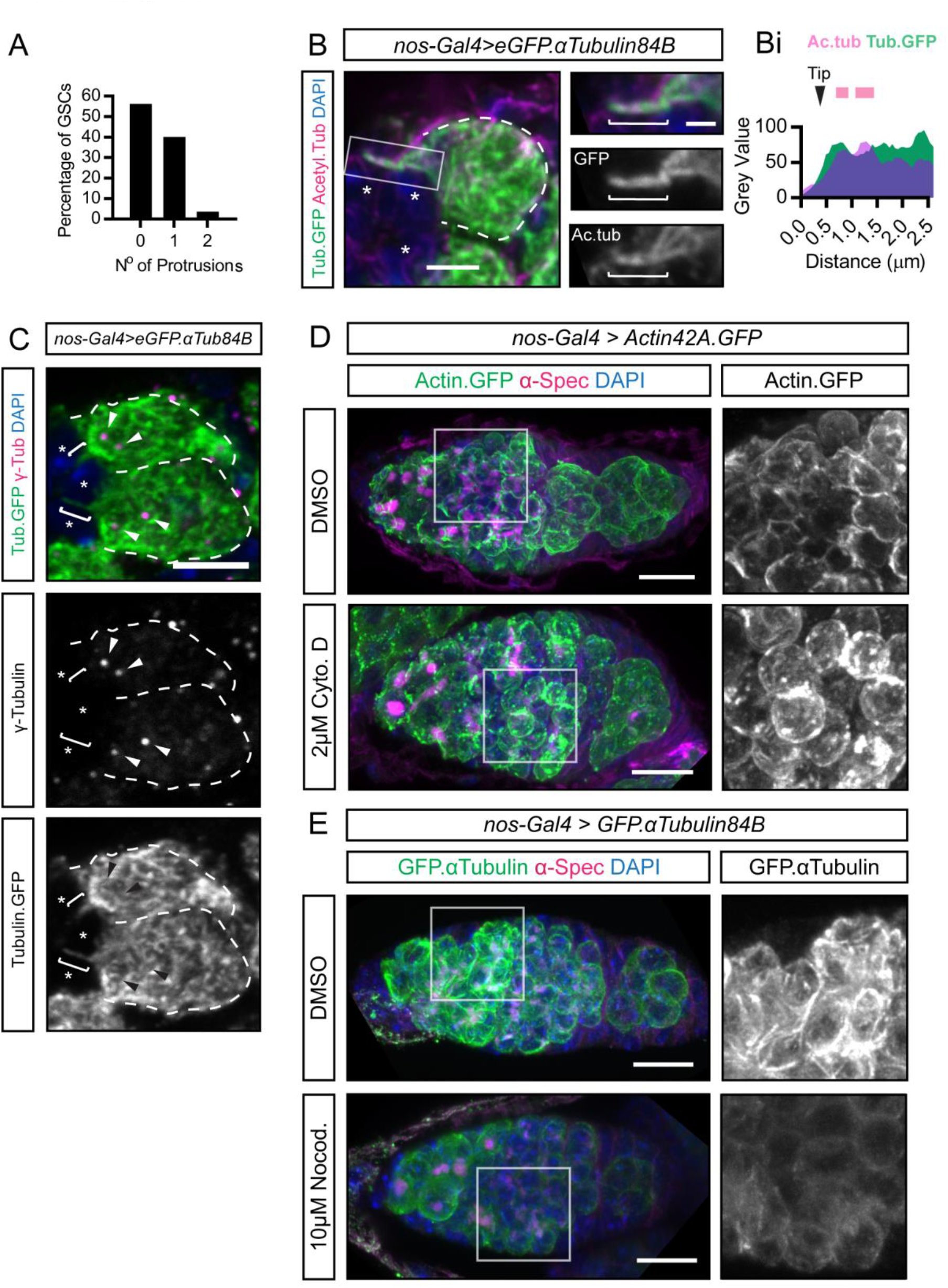
Composition of GSC projections. Refers to Figure 1. (A) Frequency of cytocensor formation per GSC. n=100 GSCs. (B) GFP-αTubulin84B marked MT-projection labelling acetylated MTs. (Inset) closeup view of boxed region. (Bi) fluorescence intensity plot along the shaft of the MT-projection. Magenta lines denote highly acetylated regions. (C) Centrosome localisation, labelled by γ-tubulin, relative to MT-projection. Brackets indicate MT-rich projections. Arrowheads indicate centrosomes. Dashed lines mark individual GSCs. (D-E) Testing *ex vivo* drug treatment on germaria expressing Actin42A.GFP or GFP-αTubulin84B. A 30 min treatment with 2µM cytochalasin D leads to fragmentation of actin and the accumulation of puncta (D). Similar treatment with 10µM nocodazole reduces tubulin levels (E). Scale bar = 5µm or 1µm (insets).

**Figure S3.**
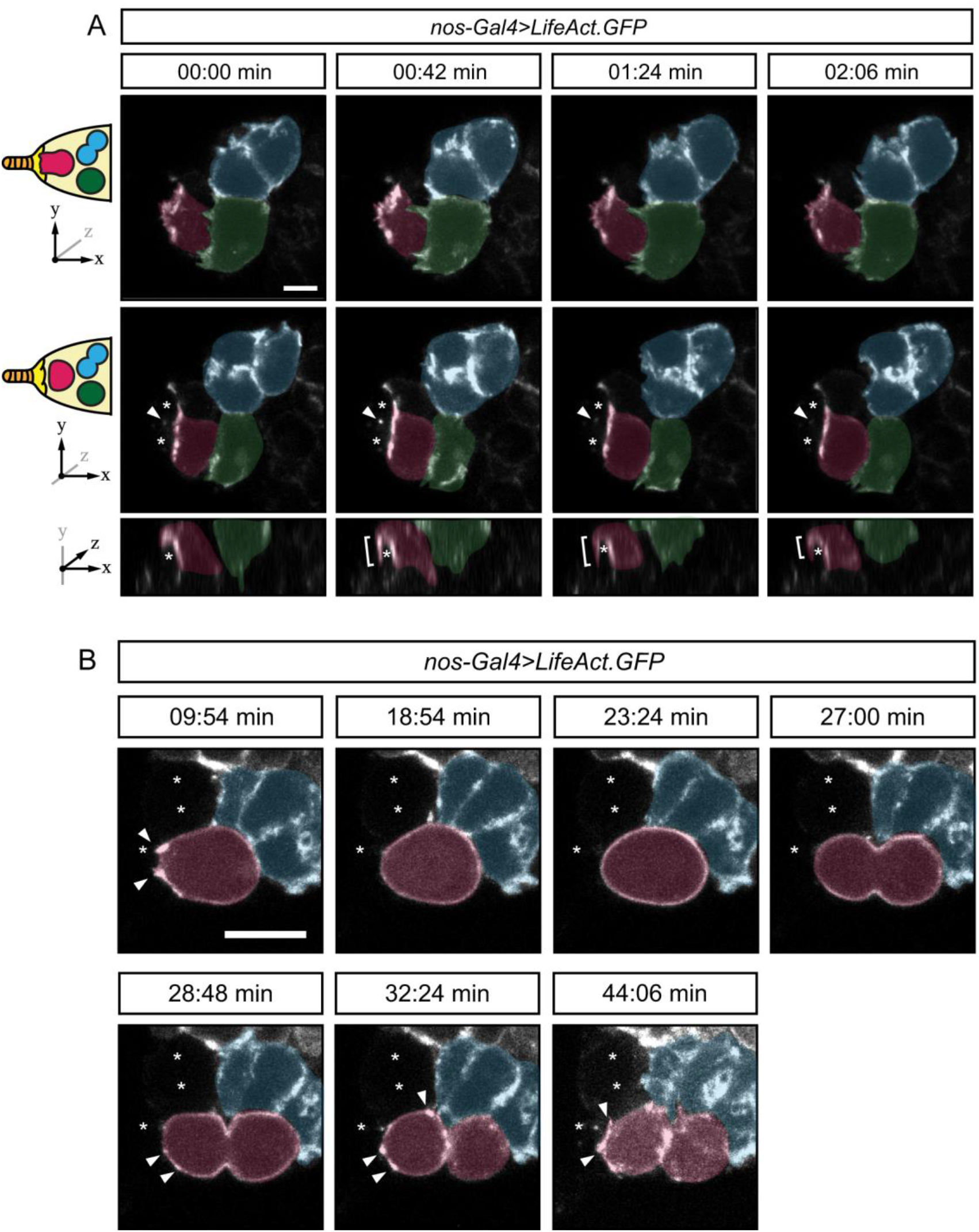
F-actin-rich projection dynamics. Refers to Figure 2. (A) Stills from Video 3 showing F-actin in GSCs. Top panels show first 2 slices of a maximum projection showing a broad lamellipodia-like projection depicted in cartoon form on the left and axes denote position within the maximum projection. Middle panels show 2 deeper slices in the middle revealing two CpCs (*) that the lamellipodial projections extend over. Bottom panel shows *xz*-plane view. Arrowheads mark the bisected shaft (middle) or brackets show the entire finger-like filpodium (bottom). (B) Stills from Video 4 showing F-actin dynamics during GSC mitosis. Arrowheads indicate F-actin-rich niche-directed projections and puncta that form post-mitosis. Niche CpCs (*). Scale bar = 5µm

**Figure S4.**
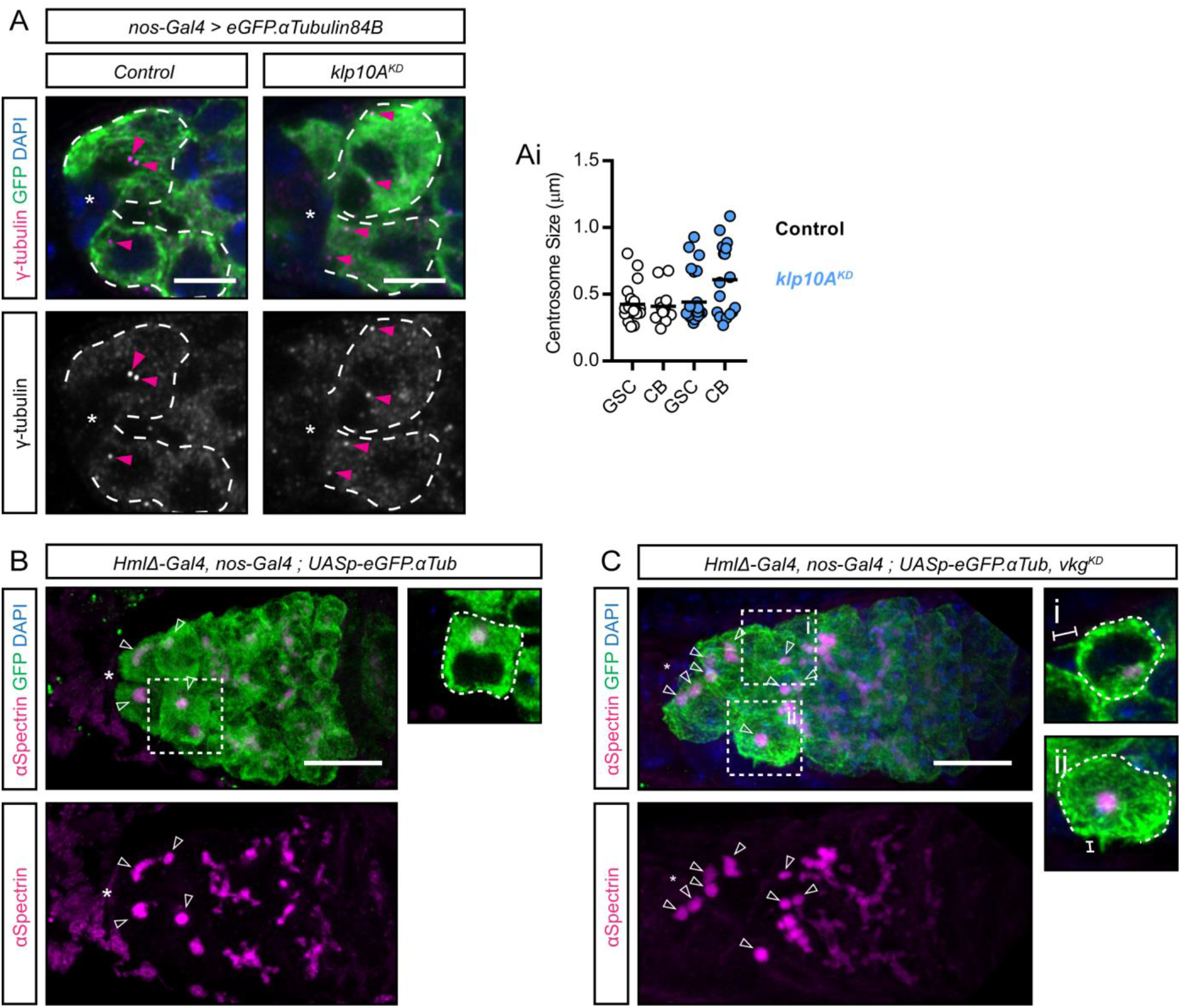
Ectopic Dpp signalling induces cytocensor formation. Refers to Figure 3. (A-Ai) Germline-specific *klp10A* knockdown has no effect on the GSC centrosome size (labelled with γ- tubulin, magenta arrowhead). (B-C) Germline and haemocyte specific expression of *eGFP-αTub84B* and RNAi-knockdown of *vkg* (ColIV) expression. (A) Control germaria have 3-4 early germ cells (open arrowheads) and cells that exit the niche (inset) do not typically generate cytocensors. (B) Knockdown of *vkg* expression in larval haemocytes extends the range of Dpp in the germarium, leading to ectopic germ cell accumulation and cytocensor formation by cells that have exited the niche (inset). Dashed lines outline individual GSCs. (*) Niche CpCs; brackets show projections. Scale bars = 5µm (A) or 10µm (B and C).

**Figure S5.**
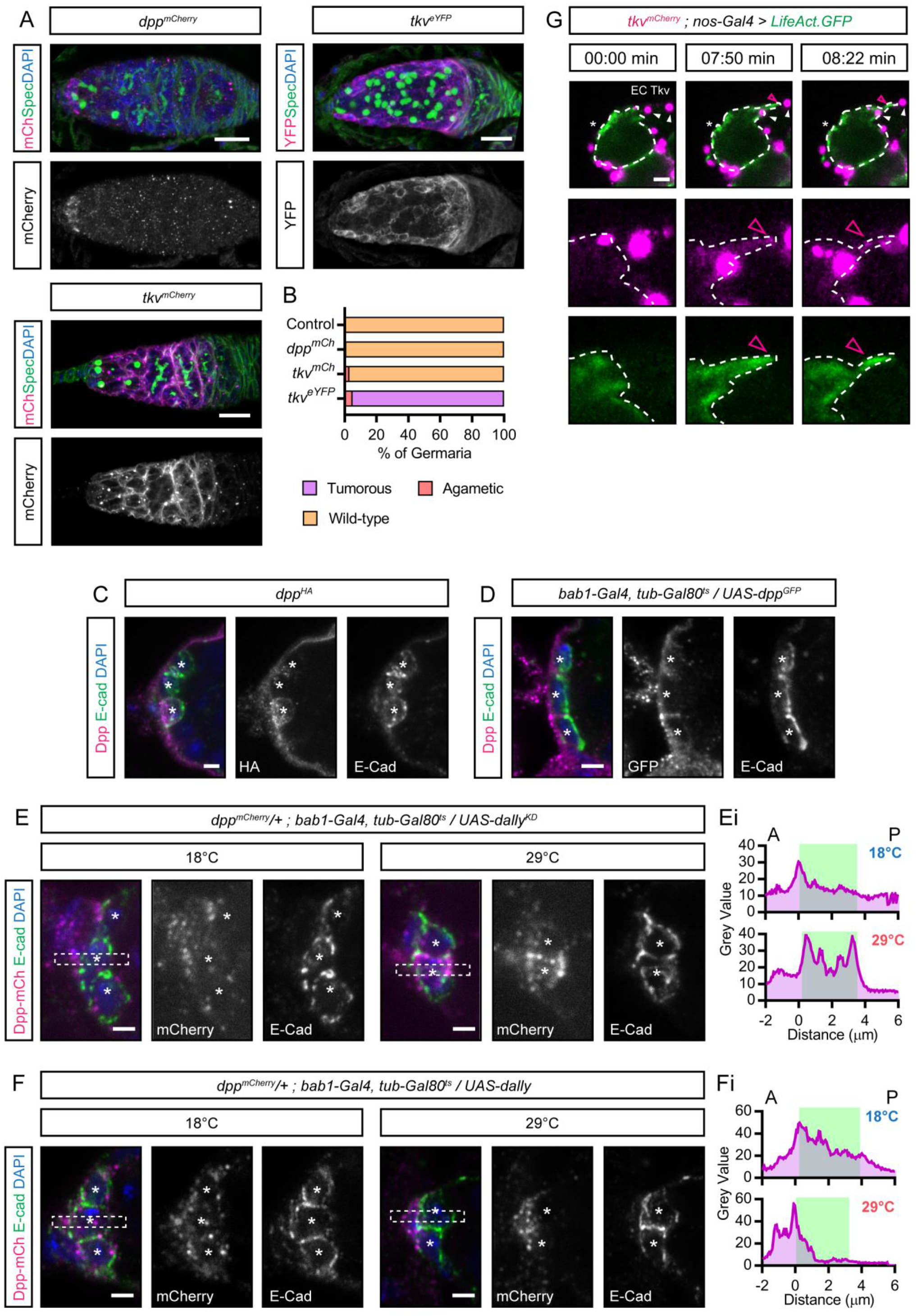
Niche cell-expressed Dally sequesters Dpp away from GSCs. Refers to Figures 4 and 5. (A) Germline phenotypic analysis of *dpp^mCh^*, *tkv^mCh^* and *tkv^eYFP^* lines quantified in (B) for n=100 germarium each. Control = wildtype. (C-D) Extracellular immunostaining of tagged-Dpp transgenic lines; (C) Dpp^HA^ under the control of its own promoter and (D) UAS-Dpp^GFP^ which was transiently expressed in the anterior somatic cells of the germarium using *bab1-Gal4, tub-Gal80^ts^*. (E-F) Dpp^mCh^ localisation around the GSC niche following transient knockdown (E) or overexpression (F) of Dally in the anterior germarial somatic cells (18°C controls and 29°C *dally* knockdown/overexpression). Box shows where the plot of fluorescence intensity (Ei and Fi) was taken from anterior to posterior (A to P) through the centre of the niche (*). E-cadherin defines the niche cell boundaries (green). (G) Stills from a video showing GSC (dashed line) F-actin labelled with *LifeAct.GFP* and endogenous mCherry-tagged Tkv showing a lateral actin-rich projection and Tkv.mCh at the tip (magenta open arrowhead). In addition, escort cell expressed ‘decoy’ Tkv are readily apparent in large puncta (white arrowhead) which are not connected to the GSC in the first panels. (*) Niche CpCs; Scale bars = 2µm.

**Figure S6.**
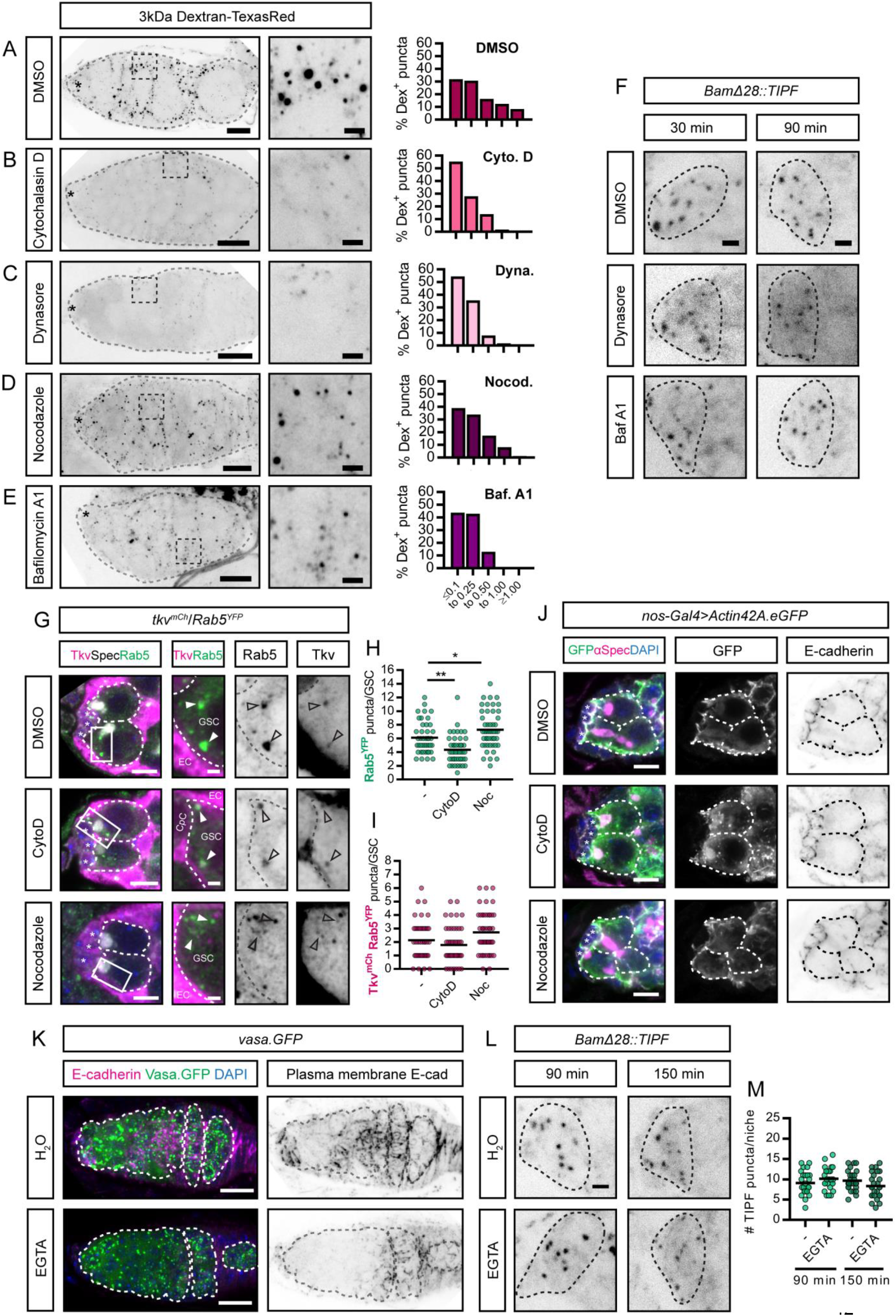
Short-term inhibition of Tkv endocytosis or degradation or niche adhesion does not affect Dpp signal transduction. Refers to Figure 7. (A-E) Endogenous fluorescence of 3kDa Dextran-Texas Red in *vasa.GFP* expressing germaria (Vasa.GFP expression not shown but used for orientation when imaging). (Insets) Closeup views of the indicated boxed regions. Histograms show the Dextran labelled puncta size as a percentage of the total number of puncta for DMSO (n=416), CytoD (n=114), dynasore (n=112), nocodazole (n=241) and BafA1 (n=240) from 5 *z-*slices taken of the entire germarium at 1µm intervals. Grey dashed line outlines the entire germarium. (F) Endogenous fluorescence of the TIPF reporter following 30 and 90 min treatments with DMSO (control), 100μM dynasore or 100nM BafA1 in inverted black and white for clarity. Black dashed lines outlines the niche CpCs. (G) Immunofluorescence staining of *tkv^mCh^ and Rab5^YFP^* expressing germaria following *ex vivo* incubation with DMSO (control), 2µM CytoD or 10µM nocodazole for 90 mins before fixation. (Insets) Closeup views of the indicated boxed regions. Individual channels are in inverted black and white for clarity. Dashed lines outline GSCs. Arrowheads mark Tkv^mCh^-positive Rab5^+^ vesicles. (H-I) Quantification of the total number of Rab5^+^ vesicles per GSC (H) and the number of Tkv^mCh^-positive Rab5^+^ vesicles per GSC (I). n ≤ 44 GSCs per treatment. (J) Immunofluorescence staining of germaria with germline-specific expression of *Actin42A.eGFP* following *ex vivo* incubation with DMSO (control), 2µM CytoD or 10µM nocodazole for 90 mins before fixation. Dashed lines outline GSCs and GSC-pCB pairs. Ecad staining is shown inverted for clarity. (K) Extracellular staining of Ecad localisation in *vasa.GFP* expressing germaria following *ex vivo* incubation with 6mM EGTA for 90 mins. Images show antibody fluorescence for Ecad and endogenous Vasa.GFP expression. Dashed lines outline Vasa^+^ germline. (L) Endogenous fluorescence of the TIPF reporter following 90 and 150 min incubations with water (control) or 6mM EGTA in inverted black and white for clarity. Black dashed lines outline the niche CpCs. (M) Quantification of the number of TIPF puncta per niche in (D) following water (control) or 6mM EGTA for either 90 mins (n = 25) or 150 mins (n=25 and 26, respectively). (*) niche CpCs. Scale bar = 10µm (A-E and K), 5µm (G and J) or 1µm (F, L and insets). *, p<0.05 and **, p<0.001.

**Figure S7.**
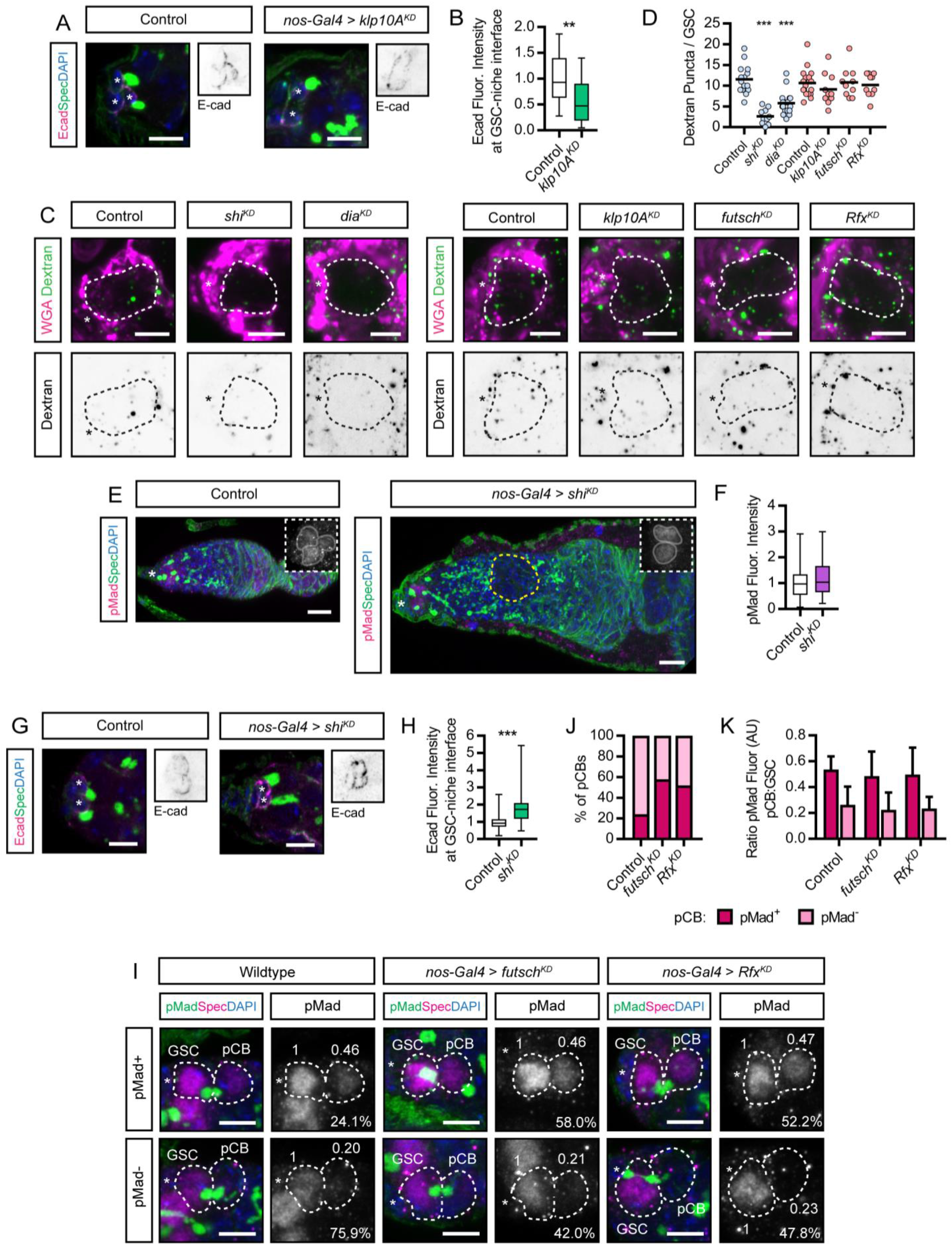
Altered endocytosis or pMad inheritance do not account for disrupted Dpp signal transduction following genetic manipulation of projection formation. Refers to Figure 7. (A-B) Immunofluorescence staining showing germline-specific *shRNA e*xpression phenotype for control and *klp10A^KD^* GSCs showing Ecad in inverted black and white for clarity and (B) quantification in control (n=31) and *klp10A^KD^* (n=21). (C) Endogenous fluorescence of 10kDa Dextran-Alexa Fluor 488 in GSCs expressing *shRNA* outlined by WGA-Alexa Fluor 633 and quantified in (D; n≥10). Blue indicates lines raised at 18°C and shifted to 25°C upon eclosion to induce RNAi only during adulthood. Red indicates lines raised at 25°C and shifted to 29°C upon eclosion to enhance RNAi. (E-F) Immunofluorescence staining showing germline-specific *shRNA e*xpression phenotype for *shi^KD^.* Early germ cells are marked by the presence of the spectrosome labelled by anti-αSpectrin. (Insets) pMad staining reports the Dpp signaling response and (F) quantification in control (n=53) and *shi^KD^* (n=54). (G-H) same as in (E) showing Ecad in inverted black and white for clarity and (H) quantification in control (n=53) and *shi^KD^* (n=55). (I) Immunofluorescence staining showing germline-specific *shRNA e*xpression phenotypes for *futsch* and *Rfx*. pMad staining reports the Dpp signalling response in GSC-pCB pairs (dashed line) identified by the shared spectrosome labelled by anti-αSpectrin. Numbers indicate the pMad levels relative to the indicated GSC and the percentage of GSC-pCB pairs that possess a pMad^+^ or pMad^-^ pCB. (J-K) Quantification of (I) showing the percentage of GSC-pCB pairs that possess a pMad^+^ or pMad^-^ pCB (J) and showing the average pMad fluorescence ratio between the GSC-pCB (K) within the pMad^+^ or pMad^-^ groups as indicated in (J). (*) niche CpCs. Scale bar = 10µm (E) and 5µm (A, C, G and I). For box and whisker graphs the box shows median, 25^th^ and 75^th^ percentile and whiskers show minima and maxima. **, p<0.001; ***, p<0.0001.

### SUPPLEMENTARY TABLES AND VIDEOS

**Table S1** Genes differentially expressed between Tkv^QD^ and bam^GFP^-expressing germ cells. Related to Figure 1.

**Table S2** Genes differentially expressed between Tkv^QD^ and bam^KD^-expressing germ cells. Related to Figure 1.

**Video S1** A dynamic cytocensor projecting into the niche. Germline expression of *UASp-eGFP.αTub*. Video represents a maximum projection over 6µm. Time = min:sec at 5 frames per second. Scale bar = 5µm. Related to Figure 2.

**Video S2** Dynamic actin-based filopodia probe the niche. Germline expression of *UASp-LifeAct.GFP*. Video represents a maximum projection over 6.75µm. Time = min:sec at 5 frames per second. Scale bar = 5µm. Related to Figure 2.

**Video S3** All early germ cells form dynamic actin projections. Germline expression *UASp-LifeAct.GFP*. Video represents a maximum projection over 3 µm. Time = min:sec at 5 frames per second. Scale bar = 5µm. Related to Figure 2 and S2.

**Video S4** Looking down the shaft of a filopodium extending into the niche from a lamellipodium. Germline expression of *UASp-LifeAct.GFP*. Video represents a maximum projection over 1.5µm. Time = min:sec at 5 frames per second. Scale bar = 5µm. Related to Figure 2 and S2.

**Video S5** GSC projections collapse prior to mitosis and rapidly reform after cell division. Germline expression of *UASp-LifeAct.GFP*. Video represents a maximum projection over 1.5µm. Time = min:sec at 5 frames per second. Scale bar = 5µm. Related to Figure 2 and S2.

